# Evaluating the Influence of Anatomical Accuracy and Electrode Positions on EEG Forward Solutions

**DOI:** 10.1101/2022.09.01.505675

**Authors:** Jesper Duemose Nielsen, Oula Puonti, Rong Xue, Axel Thielscher, Kristoffer Hougaard Madsen

## Abstract

Generating realistic volume conductor models for forward calculations in electroencephalography (EEG) is not trivial and several factors contribute to the accuracy of such models, two of which are its anatomical accuracy and the accuracy with which electrode positions are known. Here, we investigate effects of anatomical accuracy by comparing forward solutions from SimNIBS, a tool which allows state-of-the-art anatomical modeling, with well-established pipelines in MNE-Python and FieldTrip. We also compare different ways of specifying electrode locations when digitized positions are not available such as transformation of measured positions from standard space and transformation of a manufacturer layout.

Substantial effects of anatomical accuracy were seen throughout the entire brain both in terms of field topography and magnitude with SimNIBS generally being more accurate than the pipelines in MNE-Python and FieldTrip. Topographic and magnitude effects were particularly pronounced for MNE-Python which uses a three-layer boundary element method (BEM) model. We attribute these mainly to differences in skull and cerebrospinal fluid (CSF) modeling. Effects of electrode specification method were evident in occipital and posterior areas when using a transformed manufacturer layout whereas transforming measured positions from standard space generally resulted in smaller errors.

We suggest modeling the anatomy of the volume conductor as accurately possible and we hope to facilitate this by making it easy to export simulations from SimNIBS to MNE-Python and FieldTrip for further analysis. Likewise, if digitized electrode positions are not available, a set of measured positions on a standard head template may be preferable to those specified by the manufacturer.

## 1. Introduction

Electroencephalography (EEG) can be used to analyze brain activity with high temporal resolution. The measurements consist of potential differences observed on the scalp which directly reflect the synchronous activity of a large body of pyramidal cells oriented perpendicular to the cortical surface (Kirschstein and Köhling, 2009). However, the relationship between measurements and brain activity is complicated by the fact that the former is a spatially low-pass filtered representation of the latter where the filter consists of the tissues separating the neural sources from the sensors (Buzsáki et al., 2012). Due to volume conduction, all sources in the brain affect the potential at all electrodes simultaneously.

In some situations, it may be beneficial to analyze the data in the source domain rather than the sensor domain, e.g., when investigating the connectivity between different brain regions (Schoffelen and Gross, 2009; Mahjoory et al., 2017; Nguyen-Danse et al., 2021) or when localizing spike activity in epilepsy (Kaiboriboon et al., 2012; van Mierlo et al., 2020). Reconstructing the neural generators of an observed EEG signal is an inverse problem. In order to solve this, one first needs to solve the corresponding forward problem which consists of estimating the potential distribution on the scalp due to a neural source placed arbitrarily in the brain.

To solve the forward problem in EEG, one typically starts by constructing a physical model of the head, the volume conductor model, which enables simulation of volume conduction effects due to activity in the brain. Naturally, one would expect the accuracy of this volume conductor model to impact the accuracy with which such effects can be modeled (Vorwerk et al., 2014).

In this work, we investigate some of the elements which affect the accuracy of the forward solution, specifically, the anatomical accuracy of the head model and the accuracy with which electrode positions are known. We compare existing methods with SimNIBS which is able to generate state-of-the-art head models automatically based on magnetic resonance imaging (MRI) scans. Our aim is to explore the extent to which more accurate modeling leads to improvements in the forward solution and perhaps also in source localization accuracy (although the latter is not investigated here).

Numerous methods have been employed to solve the EEG forward problem. In simple geometries, e.g., nested sphere models, (quasi-)analytical solutions can be derived, however, for realistically shaped models numerical methods are needed. In the boundary element method (BEM), it is common to use a three-layer surface model (consisting of inner skull, outer skull, and skin) of relatively low resolution due to the dense nature of the problem^1^ whereas high-resolution, multi-compartment models are often seen in the finite element method (FEM). However, the sparsity of the FEM system means that it is generally inferior to BEM in terms of numerical accuracy for geometries where BEM is applicable (Hallez et al., 2007; Vorwerk et al., 2012).

One important aspect of such a model is its anatomical accuracy. In a systematic evaluation of the effect of distinguishing different tissue compartments, Vorwerk et al. (2014) concluded that modeling cerebrospinal fluid (CSF) is important as well as distinguishing white matter and gray matter, however, explicit modeling of spongy bone as well as the anisotropic conductivity of white matter were found to be less important. Using a similar strategy, Azizollahi et al. (2016) corroborated these results in neonates.

The skull compartment has received special attention in the literature due to its importance in shaping the observed fields in EEG (Hämäläinen et al., 1993). Lanfer et al. (2012) investigated the effect of several geometrical simplifications and errors on the forward solution, concluding that localized modeling errors (e.g., skull holes, erroneous thickness) result in forward errors mostly in the vicinity of such geometrical inaccuracies whereas simplifications of a more general nature (e.g., cutting the model at the base of the skull or approximating the base of the skull with constant thickness) show increased errors for large array of positions. Similar findings were reported by Chauveau et al. (2004) and Li et al. (2007). Forward modeling errors have also been observed when not modeling skull openings (Fiederer et al., 2016). Errors in skull thickness were also investigated by Chauveau et al. (2004), who found sources close to the skull to be most affected, however, their results also demonstrated that errors were seen predominantly on magnitudes and not topography. Several of these studies also report increased dipole localization errors in the vicinity of such forward modeling errors (Chauveau et al., 2004; Fiederer et al., 2016; Lanfer et al., 2012) suggesting that these effects indeed translate to errors in source localization in EEG.

To construct anatomically accurate volume conductor models, images from one or more structural sequences (e.g., T1- and T2-weighted MRI or in rare cases even computed tomography (CT)) are used. However, they may not always be available. To reconstruct sources without such anatomical information, one will have to rely on some kind of average anatomy. This was investigated by Acar and Makeig (2013) who compared individualized head models to models based on template anatomy and found median source localization errors of about 5 mm, particularly in the inferior part of the brain, since the template model was cut at the base of the skull.

Taken together, it seems that the anatomical accuracy of the forward model plays an important role in shaping not only the forward solution but likely also the final source reconstruction. FEM allows us to specify geometrically complex models, however, realistic modeling of the head geometry is difficult and has so far not been easily accessible to the EEG community. SimNIBS (Thielscher et al., 2015) is a tool for simulating electrical fields in the brain due to noninvasive brain stimulation which can construct realistic anatomical volume conductor models of the human head with reasonable accuracy from MRI scans as validated against manual segmentations based on MRI and CT (Nielsen et al., 2018; Puonti et al., 2020). Given the intimate relationship between the EEG forward problem and the problem of simulating the effect of transcranial electrical stimulation (TES)—related by reciprocity (Wolters et al., 2004; Ruffini, 2015)—SimNIBS can also be used to solve the EEG forward problem and we explore this in further detail in the current work.

Another important aspect of the forward model is how electrode positions are estimated and coregistered to the anatomical model. Different systems to acquire electrode positions exist (e.g., the Polhemus Fastrak system, https://polhemus.com), however, if such systems are not available, one may have to use a template description of the electrode positions. To this end, EEG cap manufacturers typically provide spherical angles of the different electrodes which can then be mapped onto a sphere of a radius corresponding to the size of the subject head.

Dalal et al. (2014) found that localization accuracy and output signal-to-noise ratio (SNR) of a linearly constrained minimum variance beamformer was impacted by electrode digitization technique and coregistration method. Another study by Wang and Gotman (2001) found only minimal effect of electrode position errors while simulating displacements individually for each electrode. This may be a realistic model of pure measurement errors in a scenario where positions are digitized. However, errors due to coregistration between MRI scans and digitized electrode positions or the use of template positions, might be expected to be more spatially correlated, thus introducing a general bias in electrode positions (e.g., due to tilting, stretching etc.) (Acar and Makeig, 2013; Homölle and Oostenveld, 2019). Homölle and Oostenveld (2019) showed that digitizing electrodes using a structured-light 3D scanner resulted in slightly smaller errors compared to transformation of a custom template which again performed better than simply using the manufacturer template positions. Errors were also evident in the forward solution and a subsequent dipole fit. The errors of the latter reached 30 mm over the parietal cortex for the manufacturer template coinciding with the largest spatial bias in electrode positions. Acar and Makeig (2013) simulated coregistration errors by tilting electrode positions and found increased source localization errors (on average 5 mm) affecting predominantly superficial sources. Thus, we expect the accuracy with which electrode positions are determined to affect not only the forward model but also subsequent source estimates.

Here we explicitly consider how these two aspects of the forward model, namely its anatomical accuracy as well as how electrode positions are estimated, affect the forward calculations. The geometrical specification of the model will, to a large extent, be determined by the particular choice of forward modeling pipeline. Consequently, we first compare pipelines from different software packages for generating such models. This will result in aggregated effects from various sources but also provide practically relevant insights into the performance of the tested pipelines. Specification of electrode locations will depend on the available equipment and resources. Since it may not always be possible to obtain individual estimates during the experimental session, one may have to resort to a set of template positions. Thus, we compare different template layouts to determine their effect on the final solution in the second part of the study.

Finally, although not the focus here, we would like to point out that several other parameters of the forward model are important as well, such as the influence of the conductivity profile of different tissues. The conductivity of some tissues in the human head are fairly well known (e.g., that of CSF) (Baumann et al., 1997) whereas that of others (e.g., compact and spongy bone) (Ümit Aydin et al., 2014) generally show large variability across studies. Even though this is known to affect modeling accuracy (Bangera et al., 2010; Dannhauer et al., 2011; Marin et al., 1998; McCann and Beltrachini, 2022; Vallaghe and Clerc, 2009; Vorwerk et al., 2019), one will typically have to resort to standard values from the literature although calibration paradigms have also been suggested with (Ümit Aydin et al., 2014) or without (Acar et al., 2016) the use of magnetoencephalography. Finally, model resolution is also important (Kaipio and Somersalo, 2007). In most cases, though, researchers will probably rely on the default settings of a particular pipeline (as determined from validation experiments).

We start by evaluating the numerical accuracy of SimNIBS in a spherical model and a realistic three-layer head model. Next, we compare forward solutions from SimNIBS with standard pipelines in MNE-Python and FieldTrip. We also include a model based on standard anatomy representing a scenario where electrode positions are known but the anatomy is not. As we will see, the pipelines differ substantially in terms of anatomical accuracy. Finally, we compare different ways of specifying electrode positions when digitized positions are not available such as transformation of positions from standard space and transformation of manufacturer layout positions. As in the first sub-study, we also include a model based on standard anatomy.

## 2. Methods

### 2.1. Calculating EEG Forward Solutions in SimNIBS

Starting from the quasi-static approximation of the Maxwell equations, the EEG forward problem takes the form

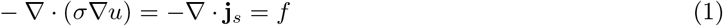

subject to boundary conditions

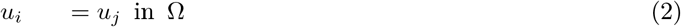

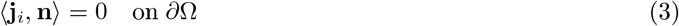

where *i* and *j* are two neighboring elements (e.g., tetrahedra) and **n** is the vector normal to their interface. 〈**j**, **n**〉 is the projection of the current density, **j**, in the normal direction, and Ω and *∂*Ω refer to the interior of the domain and its surface, respectively. Equation (2) ensures that the solution is continuous across interfaces of neighboring elements whereas equation (3) states that no current can leave the domain. We are interested in finding *u,* the potential distribution, for a given *f* (assumed to be known), which can be interpreted as a monopolar source configuration, i.e., the source of the current density, **j**_*s*_ (Hallez et al., 2007).

Equation (1) states that the divergence of the return current density (left-hand side) is equal to the divergence of the source current density. This is important for EEG source analysis as we measure potential differences on the scalp due to the return currents induced by the primary currents (i.e., the neural sources) along with other effects of non-interest (e.g., respiration, heartbeat).

Due to the principle of reciprocity, the EEG forward problem is related to the estimation of electric fields induced by noninvasive brain stimulation. Specifically, the potential between two points, *a* and *b,* due to a dipolar source at a certain position is related to the electric field at that position resulting from running a current between *a* and *b* (Weinstein et al., 2000). SimNIBS is a software package for solving the latter problem. To generate so-called leadfields for EEG, we exploit the intimate relationship between the EEG forward problem (estimating scalp potentials from current sources in the brain) and TES (estimating the electric field resulting from applying a potential difference between two electrodes or, equivalently, applying a current at one electrode and removing it at another). SimNIBS returns a smooth estimate of the electric field from which the gain (or leadfield) matrix can be computed. We describe how this is achieved below.

In SimNIBS, the volume conductor model is constructed using tetrahedral elements (potentially with anisotropic conductivities) and linear basis functions are employed to model the potential. The electric field is estimated by differentiating the solution and nodal values are recovered separately for each tissue compartment using superconvergent patch recovery (SPR) (Zienkiewicz and Zhu, 1992). In EEG, the sources are located in the gray matter and we therefore interpolate the field on a surface representing the center of the gray matter sheet.

Having solved the equation system for each electrode wrt. the reference, we start by computing the (negative) gradient of the solution to obtain the electric field, **e**_*i*_, in each element, *i*,

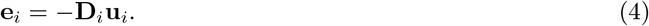

where **D**_*i*_ is the gradient operator and **u**_*i*_ contains the potentials at the nodes of *i*. As we use linear basis functions, this field estimate is not continuous and therefore very inaccurate for locations other than the element barycenters. To recover nodal field values, we use SPR to construct a smooth, interpolating function for each node by fitting a linear model to the values in an element patch (i.e., the collection of all elements associated with a given node) around each node such that a field component at an arbitrary position, *r*, can be estimated as

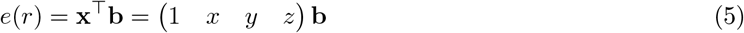

where (*x, y, z*) are the coordinates at position *r* and **b** is a vector of parameters to be estimated. This is solved by minimizing the least squares error between the values of the function **f** which we are trying to fit (here the values of the electric field) and the interpolant at the barycenters

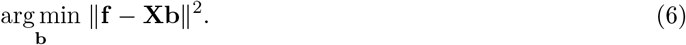

Here, **X** is constructed by row-wise stacking of components 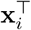 corresponding to the barycenter of each element in the patch (augmented by a column of ones as above). We can compute the projection matrix at *r* as **x**^⊤^ (**X**^⊤^**X**)**X**^⊤^ and apply it to each component of the electric field to interpolate the field at this location (which, in this case, is the coordinates of the node defining the element patch). Finally, the electric field at any location can be estimated using barycentric interpolation (see supplementary material of Saturnino et al., 2019, for details).

Following Ruffini (2015)^2^, if we consider a particular location in the brain, *r*, and a pair of electrodes, *a* and *b,* on the scalp, we have

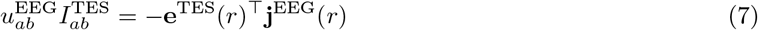

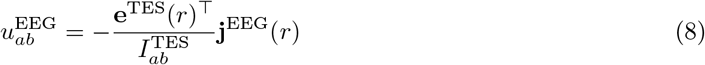

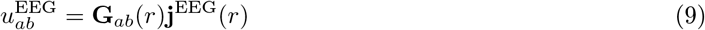

where **e**(*r*) is the electric field induced by TES at position, *r*, using current *I*, and *u* is the potential difference at *a* wrt. *b* due to a dipole source **j**(*r*). **G**_*ab*_(*r*) is the potential field value at *a* wrt. *b* for unit dipoles at *r* aligned with each principal axis. Collecting these for all locations and electrode pairs and augmenting it with the reference electrode (effectively, adding zeros corresponding to the reference electrode, and applying an average reference), we arrive at the final matrix of leadfields, i.e., a matrix describing potential fields induced by (ideal) dipolar sources in the brain. Efficient implementations based on reciprocity have also been presented by others (Weinstein et al., 2000; Wolters et al., 2004).

### 2.2. Validation of SimNIBS for EEG

We validate the leadfields generated using SimNIBS in two ways. First, we compare with an analytical solution in a simple geometry consisting of nested spheres. Second, using a realistically shaped three-compartment model, we compare with existing BEM implementations as BEM generally has high numerical accuracy compared to first order FEM.

#### 2.2.1. Spherical Model

We constructed a model consisting of three concentric spheres representing the inner skull, outer skull, and outer skin surfaces with radii 80, 86, and 92 mm and conductivities 0.3, 0.006, 0.3 Sm^-1^, respectively. We generated such models with different node densities, 0.065, 0.125, 0.25, 0.5, and 1.0 nodes/mm^2^ (see table 1 for information on number of nodes and elements in the volume meshes), to investigate the effect of resolution on the numerical accuracy and used Gmsh (Geuzaine and Remacle, 2009) to generate a volume mesh representation of each of the spherical models. Following Vorwerk et al. (2012), we placed point sources aligned with each axis (*x, y,* and *z*) in steps of 1 mm ranging from 2 to 77 mm to the sphere origin. This was done for 1,000 random directions. Thus, all sources resided within the innermost compartment of the model. The outermost sources were at a distance of 15 mm to the skin surface which is similar to the closest sources in a realistic head model (Lu et al., 2019). Finally, we placed 100 electrodes equidistantly on the outermost surface. In the simulations, these were modeled as rings with 10 mm radius and 4 mm thickness and were meshed onto the spherical models whereas in the analytical solutions they were treated as point electrodes^3^. The conductivity of the electrodes was set to 29.4 Sm^-1^. Analytical potentials were computed at the location of the electrodes using equations 2 and 2a from Ary et al. (1981).

**Table 1.** Resolution of surface meshes and number of nodes and elements in the corresponding volume meshes used in the validation experiments. The number and “C” in parenthesis refers to the number of compartments in the model.

| surface<br>nodes/mm<sup>2</sup> | volume |  |
| --- | --- | --- |
|  | nodes | tetrahedra |
| spherical models (3C) |  |  |
| 0.065 | 24 880 | 126 844 |
| 0.125 | 61 154 | 322 872 |
| 0.25 | 133 952 | 710 780 |
| 0.5 | 356 742 | 1 934 983 |
| 1.0 | 903 074 | 4 977 163 |
| realistic models (3C) |  |  |
| low | 22 969 | 120 130 |
| high | 697 793 | 3 861 891 |
| realistic models (5C) |  |  |
| 0.5 | 638 247 | 3 530 439 |
| 0.5 (refined) | 4 880 956 | 28 243 512 |
| 1.0 | 1 955 900 | 11 171 938 |
| 1.0 (refined) | 15 228 577 | 89 375 504 |

#### 2.2.2. Realistic Geometry

To validate the models in a realistically shaped geometry, we extracted surfaces corresponding to inner skull, outer skull, and skin of a head model generated using the *headreco* pipeline in SimNIBS. We downsampled each surface to approximately 5,000 nodes and used these in the simulations. Additionally, we upsampled these surfaces by uniform remeshing using MeshFix (Attene, 2010) to a resolution close to what we would normally use in our FEM calculations (approximately 80,000 nodes). Running this model in BEM was not feasible with the implementations used here due to the excessive computational resources needed. Conductivities were the same as in the three-layer sphere model. Electrodes were modeled as described above and were placed by creating a standard montage of the 64 channel BioSemi cap (https://www.biosemi.com) included in MNE-Python. For comparison, we used BEM implementations from FieldTrip (specifically, the *dipoli* implementation) and MNE-Python, the latter of which was used as a reference.

We also show numerical differences between FEM models with different surface densities (specifically, 0.5 and 1.0 nodes/mm^2^) and refined (achieved by splitting the tetrahedra) versions of these models to illustrate the effect of geometrical accuracy compared to that of resolution. See table 1 for information on number of nodes and elements in the volume meshes.

Lastly, we show results of a comparison between a five-compartment FEM model and a three-layer BEM model. This allows us to compare the numerical differences between FEM and BEM (from the previous simulation) with the differences resulting from modeling the anatomy at different levels of geometrical accuracy and detail. Here, the FEM model was used as reference.

### 2.3. Impact of Anatomical Accuracy on the Forward Solution

The dataset used in this study consists of 20 subjects each of which had a T1- and a T2-weighted MRI scan (for details on this dataset, please see Farcito et al., 2019). Importantly, manual segmentations of 16 tissue classes were available for all subjects. However, in this study we only distinguished 9 tissues^4^ (10 including air pockets).

#### 2.3.1. Volume Conductor Models

To study the effect of anatomical accuracy we generated four head models using different approaches, specifically, (1) SimNIBS with complete head anatomy reconstruction method (CHARM) (a new segmentation and meshing pipeline which is available in SimNIBS 4.0) (Puonti et al., 2020), (2) MNE-Python (Gramfort, 2013) which is based on FreeSurfer (Dale et al., 1999), (3) FieldTrip (Oostenveld et al., 2011) which uses SPM12 (https://www.fil.ion.ucl.ac.uk/spm/software/spm12/) for segmentation and DUNEuro (Schrader et al., 2021) for FEM modeling, and (4) SimNIBS with CHARM using standard anatomy. As a reference, we used the manual segmentations which we meshed using SimNIBS^5^. Here we briefly describe each of the pipelines.

##### SimNIBS-CHARM

The CHARM pipeline uses the SAMSEG framework (Puonti et al., 2016) to segment one or more MRI scans into fifteen different tissue classes. Subsequently, the segmentation is upsampled to 0.5 mm^3^ isotropic resolution and post-processed using morphological operations to smooth the segmentation and to ensure that the brain is surrounded by at least one voxel (0.5 mm) thick layer of CSF. The central cortical surface is reconstructed using adapted versions of the corresponding CAT12 (Dahnke et al., 2013) functionalities. The pial surface is then estimated from the central cortical surface and used to improve the sulci representations in the original segmentation. Finally, a volume mesh representation is generated using GGAL (https://www.cgal.org) by directly meshing the updated segmentation. The size of the tetrahedral elements are locally adapted using sizing fields such that elements that are far from tissue borders are larger and elements that are close to tissue borders are smaller. The initial tetrahedral mesh is post-processed to improve the quality of the tissue surfaces. This is done by reassigning the tissue type of tetrahedra representing “spikes” in the surfaces. In addition, the surface nodes are smoothed using a Taubin approach (Taubin, 1995) which is adapted to ensure good quality of the tetrahedra connected to the surfaces. These steps are implemented as standard steps of the CHARM segmentation and meshing pipeline.

Please note that the dataset described in section 2.3 was used to create the tissue priors used in CHARM (Puonti et al., 2020), however, when running CHARM, we used priors from a four-fold cross-validation split such that a subject was never included in the prior used when creating its segmentation to circumvent biased accuracy estimates due to overfitting. The FEM simulations were run using standard settings except that the Pardiso FEM solver included in the Intel Math Kernel Library (version 2022.0.1) was used to calculate multiple FEM solutions efficiently.

##### MNE-FS

MNE-Python integrates with FreeSurfer and relies on its *recon-all* pipeline. Here we created the BEM surfaces (inner skull, outer skull, and skin) based only on a T1-weighted image^6^. This procedure uses the watershed algorithm in FreeSurfer. The final node count in each surface was 5,120. We note that since the inner and outer skull surfaces sometimes overlapped, we had to fix this in order to be able to run the BEM calculations. This was done using MeshFix by pushing the inner skull inwards to ensure a minimum distance of 2 mm between the inner and outer skull since the problem seemed to be that the inner skull surface extended too far outwards. Consequently, we view this as an improvement of the model. We used MNE-Python 0.24 and FreeSurfer 7.1.0.

##### FieldTrip-SPM

FieldTrip^7^ uses SPM12 to segment a T1-weighted image into five tissue classes (gray matter, white matter, CSF, bone, and skin). This segmentation is post-processed using morphological operations (e.g., smoothing) to, for example, ensure that the brain is surrounded by CSF which again is surrounded by bone. Based on this segmentation, a hexahedral mesh was created with a shift of 0.3 and a downsampling factor of two. We used the DUNEuro solver with default settings.

##### MNI-Template

This model is created simply by running CHARM on the MNI152 T1-weighted image. We padded the image such that the lower part of the face and neck was filled from the prior. Otherwise the procedure is exactly as described for SimNIBS-CHARM.

The purpose of this work is not to provide a thorough evaluation of the anatomical accuracy of each of these models (we refer to Nielsen et al. (2018) and Puonti et al. (2020)), however, visual inspection revealed that the SimNIBS-CHARM models were generally more anatomically accurate than the other models (see figures 6 and 7 for two examples). To summarize, SimNIBS-CHARM is a first order FEM model with generally high anatomical accuracy; FieldTrip-SPM is a FEM model with decent accuracy of brain tissue but simplified non-brain tissue (e.g., bone); MNE-FS is a BEM model with simplified anatomy (no distinction between white matter, gray matter, and CSF) and limited accuracy; MNI-Template is a FEM model based on average anatomy which, in general, is quite dissimilar to the individual anatomy but may estimate CSF and bone reasonably well.

In each case, we used the default conductivities specified by the particular pipeline^8^.

#### 2.3.2. Electrode Positions

To model electrode positions we used an EasyCap BC-TMS64-X21 transcranial magnetic stimulation (TMS) compatible cap with a modified M10 (equidistant) layout (https://www.easycap.de). See section 2.4.1 for details on how these positions were obtained.

We placed the electrodes on the individual reference models (i.e., the models based on the manual segmentations) by registering the template positions (in MNI space) to subject MRI space by a seven parameter affine registration (one uniform scaling parameter) based on coregistration of landmarks (nasion, left preauricular (LPA), right preauricular (RPA)). For SimNIBS-CHARM, MNE-FS, and FieldTrip-SPM, the electrodes were then projected onto the skin surface of the respective model. For MNI-Template, the mesh was deformed nonlinearly using thin-plate splines to head points sampled around the electrodes (four points per electrode resulting in approximately 200 points in total) similar to the procedure used in Brainstorm (Darvas et al., 2006). Given that the skin surface was well modeled in all pipelines, we do not anticipate major effects due to different electrode positions.

#### 2.3.3. Source Space

In the MNE-FS and FieldTrip-SPM models, we defined the source space as the central gray matter surface extracted from the reference model using CAT12 and downsampled to approximately 10,000 nodes per hemisphere. In the event that a source was not inside the brain (defined as the brain compartment in MNE-FS and white matter or gray matter in FieldTrip-SPM) this was excluded from analysis^9^. In SimNIBS-CHARM and MNI-Template, we used the the central gray matter surface created during the respective runs. To compare the gain matrices (leadfields) we mapped them to *fsaverage* before computing the evaluation metrics.

### 2.4. Impact of Electrode Positions on the Forward Solution

The dataset used in this study is a subset of the dataset described in Madsen et al. (2019) and Karabanov et al. (2021) in which noninvasive brain stimulation was performed conditional on the brain state as determined by EEG measurements. Of particular relevance to the current study, structural MRI scans were acquired and electrode positions digitized using an infrared optic stereo tracking system from Localite (https://localite.de). For this study, we extracted 32 subjects for which T1- and T2-weighted images as well as digitized electrode positions were available. During our analysis, we found that some digitization runs contained electrode positions which were clearly the result of some kind of equipment malfunction (e.g., a situation where an electrode was placed in the center of the head). To formalize the identification of such errors, and possibly “correct” them, we first transformed all electrode positions to standard space by nonlinear deformation. We then defined a “spurious digitization” as a position which was more than five centimeters away from the group median and we decided to correct a maximum of one electrode per subject. Corrections were performed by replacing the erroneous coordinates with the group median. Spurious digitization errors were detected in four of 32 subjects. In each of these subjects there was only a single error which we then corrected.

#### 2.4.1. EEG Custom Template Generation

We generated a custom template in standard space of the EasyCap BC-TMS64-X21 EEG cap using the following procedure. We reconstructed the skin surface of an augmented MNI152 template (which included the jaw and neck as well) from the final segmentation of the MNI-Template described in section 2.3.1 using the marching cubes implementation from *scikit-image* (van der Walt et al., 2014) and smoothed the result using *PyVista* (Sullivan and Kaszynski, 2019). Since the MNI152 model is quite large (approximately 60 cm circumference of the head), we scaled it to a circumference of 57 cm which allowed us to comfortably fit a 58 cm cap on it. This scaled model was 3D printed on an Ultimaker 2 Extended+, the aforementioned cap was placed on it, and electrode positions were digitized using a Polaris Spectra infrared camera from NDI with custom software (https://www.ndigital.com). To securely fasten the sensor responsible for tracking the head position, we augmented and flattened the nasal bridge such that it could be easily attached here. The final template was scaled back to the original space of the MNI template.

To facilitate coregistration between the coordinate system of the digitizer and MNI space, we added 13 small dents (which served as “landmarks”) in the surface (nasion, LPA, RPA, and ten points spread across the top of the head). We placed the EEG cap according to standard guidelines (e.g., Homölle and Oostenveld, 2019) and digitized the electrode positions as well as the landmarks. We repeated this three times, registered the electrode positions to MNI space and used the median position for the final template. In the final template we noticed a few irregularities. First, two parietal channels, 50 and 52, were not exactly equidistant to the surrounding electrodes. Upon inspection of the different cap sizes, we found that these channels were placed on one side of a stitching in caps smaller than 56 cm whereas they were on the opposite side in caps larger than or equal to 58 cm. Since this effect was evident mostly in the lateral direction, channels 50 and 52 were corrected by replacing the *x* coordinate with the mean of electrodes (49, 51) and (51, 53), respectively. Likewise, we saw variations in channel 22 between caps. Channel 22 was corrected by replacing the *x* and *y* coordinate with the means from 21 and 23. Finally, all channels were (re)projected onto the MNI skin surface. These steps were taken to improve the average fit of the template to subjects of different head sizes.

#### 2.4.2. Volume Conductor Models

To study the effect of different ways of estimating electrode positions, we first constructed a model of the head for each subject using the CHARM pipeline in SimNIBS. To assess the errors in electrode positions when using template layouts, we transformed the layout of the EEG cap as defined by the manufacturer^10^ as well as our custom positions from standard space to subject space. The former was transformed using an affine transformation obtained by coregistering fiducials (nasion, LPA, and RPA) between subject space and the space of the manufacturer layout (after mapping the positions to a sphere). The latter was transformed in three different ways using (1) the nonlinear deformation field calculated by CHARM as part of the segmentation, (2) the affine registration between subject space and MNI space estimated by CHARM before performing the segmentation, and (3) the affine transformation obtained by coregistration of fiducials in subject space and MNI space. Subject specific fiducials were identified on the T1-weighted MRI of each subject. As reference, we used the digitized positions available in the data set.

We also included a model based on standard anatomy (similar to the one described in section 2.3.1). As in the reference model, we used the digitized electrode positions from the data set. This was done to model a scenario where electrode positions have been accurately defined but structural information is unavailable.

As we did not find any appreciable differences in the accuracy of the channel positions between different ways of transforming the custom template (see section 3.3), we chose to only use the nonlinearly deformed positions in the forward simulations as this does not require identification of landmark positions for each individual. Thus, we ran simulations using the models with the digitized positions, nonlinear deformation of custom template (Custom-Template), with affine registration of the manufacturer layout (Man-Template), and with standard anatomy (MNI-Digitized). All simulations were carried out in SimNIBS similarly to what was described for SimNIBS-CHARM in section 2.3.1.

#### 2.4.3. Source Space

As source space we used the central gray extracted using the CAT12 functionalities embedded in CHARM and downsampled to approximately 10,000 vertices per hemisphere, thus the source space was the same for models based on individual subject anatomy (the reference using the digitized positions from the data set, Custom-Template, Man-Template) but different for the one based on standard anatomy (MNI-Digitized). To compare the gain matrices we mapped them to *fsaverage* before computing the evaluation metrics.

### 2.5. Evaluation Metrics

To evaluate differences in electrode position, we used the Euclidean distances to the reference positions. We show the full distribution of these errors and also how the mean errors are distributed over channels. Additionally, we also show the spatial distribution of the absolute error along each axis (*x, y, z*) to explore the direction of the errors. For evaluation, we always used channel positions after projecting onto the skin surface.

All channel positions were evaluated in the head coordinate system used by MNE-Python. Here, the *x*-axis goes through the LPA to the RPA, the *y*-axis is normal to the *x*-axis and passes through the nasion, and the *z*-axis is normal to the *xy*-plane and points upwards according to the right-hand rule. Thus, this constitutes a right-anterior-superior (RAS) oriented coordinate system.

To evaluate the differences between forward models we use the relative difference measure (RDM)

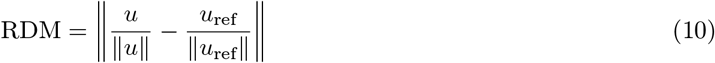

and the logarithm of the magnitude error (lnMAG)

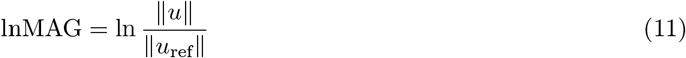

which computes the relevant measure of *u* with respect to *u*_ref_ and where ||·|| is the 2-norm (Meijs et al., 1989; Vorwerk et al., 2014). RDM measures the topographic error and is bounded between 0 and 2 where 0 means no difference whereas lnMAG measures the magnitude errors with a value of 0 corresponding to no difference.

All forward solutions were re-referenced to an average reference and converted to a fixed orientation (normal to the cortical surface). The final gain matrix consisted of approximately 20,000 columns. Before computing the evaluation metrics, gain matrices were morphed to the fifth subdivision of the *fsaverage* template *(“fsaverage5”)* which consists of 20,484 nodes in total (10,242 per hemisphere).

We show the cumulative relative frequency (CRF) of the error metrics using the data from all subjects and also the spatial distribution of the average of each metric. All surface-based plots were created using *PyVista* 0.35.1 (Sullivan and Kaszynski, 2019). Remaining plots were created with *Matplotlib* 3.5.1 (Hunter, 2007).

## 3. Results

### 3.1. Validation of SimNIBS for EEG

#### 3.1.1. Spherical Model

Figure 1 shows the CRF of RDM and lnMAG for each model and figure 2 shows RDM and lnMAG as a function of source eccentricity. Both figures illustrate that increasing the resolution of the model is beneficial for numerical accuracy. Errors get larger with increasing source eccentricity. However, for the high density models, the RDM stays below 0.02 and the lnMAG between ±0.01. From figure 2, we see a small bias in the magnitude in that it is smaller in the SimNIBS simulations.

**Figure 1.**
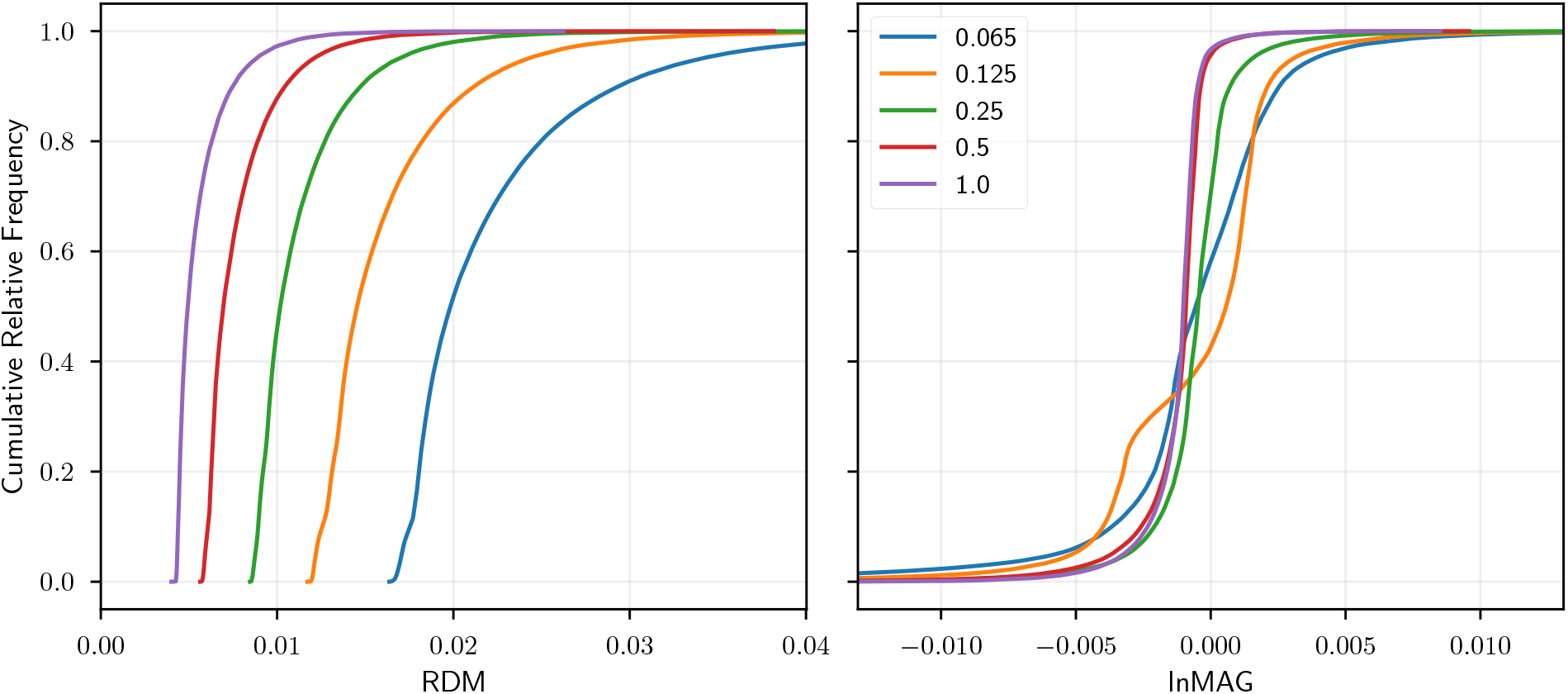
CRF of RDM and lnMAG for different model densities (nodes/mm^2^) in a three-layer sphere model.

**Figure 2.**
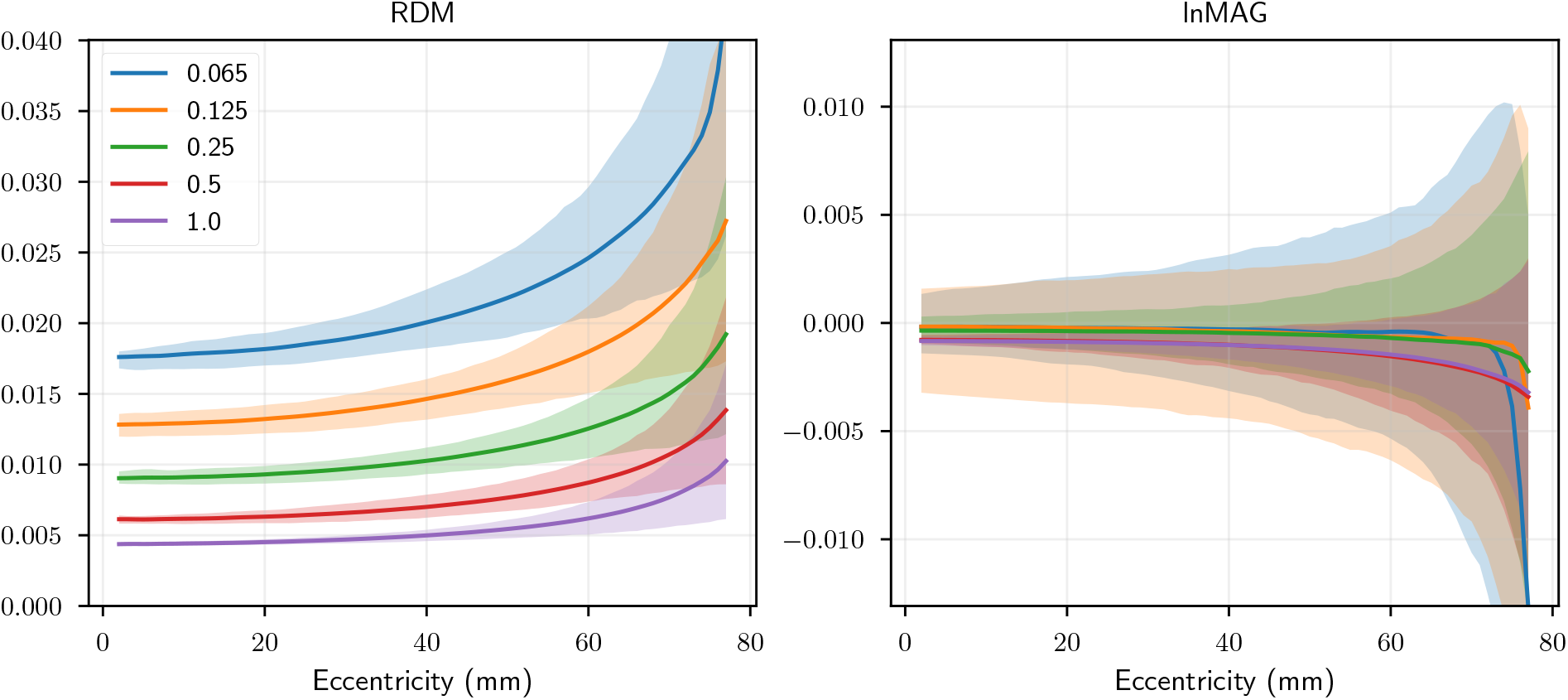
RDM and lnMAG as a function of source eccentricity for different model densities in a three-layer sphere model. Shaded area denote the interval from the 0.5th to 99.5th percentile.

#### 3.1.2. Realistic Geometry

Figure 3 shows CRF of RDM and lnMAG in a realistic three-layer model using the BEM solution from MNE-Python as reference. We see that errors of the FEM model with a resolution used in practice, i.e., the high resolution model, are generally low (below 0.05) except for a few outliers (heavy tail). The numerical accuracy generally deteriorates as sources get closer to the sensors (see section S1 and fig. S1.1 in the supplementary material). The BEM implementations are very similar in terms of topography but less so for magnitude.

**Figure 3.**
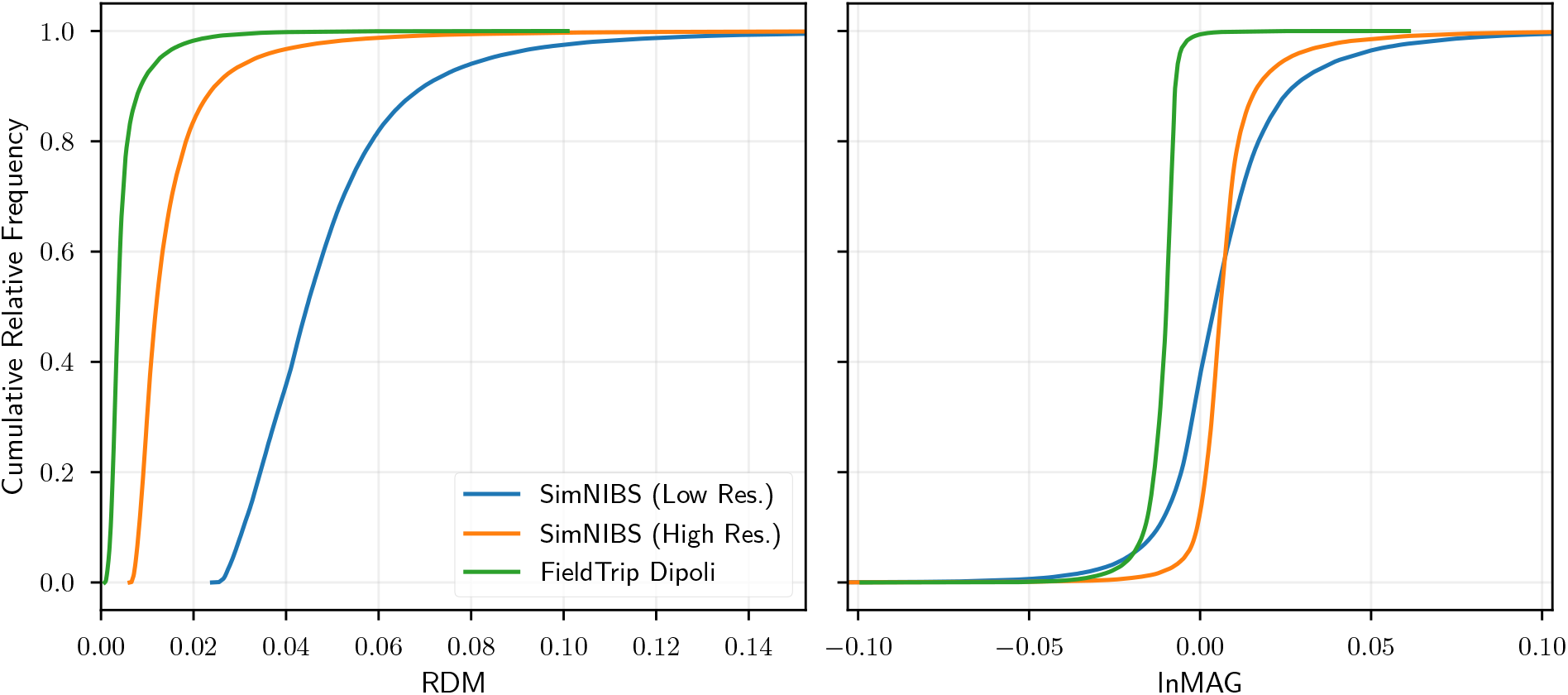
CRF of RDM and lnMAG for a high and low resolution FEM model and a BEM model in a realistically shaped three-compartment model. The BEM solution from MNE-Python was used as a reference.

Figure 4 shows CRF of RDM and lnMAG for different model comparisons. We see that purely numerical errors (as evidenced by the comparison of a model with its refined counterpart, e.g., FEM 0.5 vs FEM 0.5r) are lower than errors due to modeling the anatomy in greater detail (i.e., FEM 0.5 vs. FEM 1.0) which again is much lower than the errors incurred by simplifications of the model (i.e., the full FEM models vs. three-layer BEM). An example of the effect of resolution and refinement on the geometry of the FEM models are illustrated in figure 5. Errors due to model simplifications are about ten times those of the errors due to numerical inaccuracies. The pattern is the same for both RDM and lnMAG. Contrary to the simple three-layer model evaluated above, source eccentricity does not seem to be as good a predictor of the overall errors (numerical and otherwise) in more complicated geometries (see section S1 and figs. S1.2 and S1.3 in the supplementary material).

**Figure 4.**
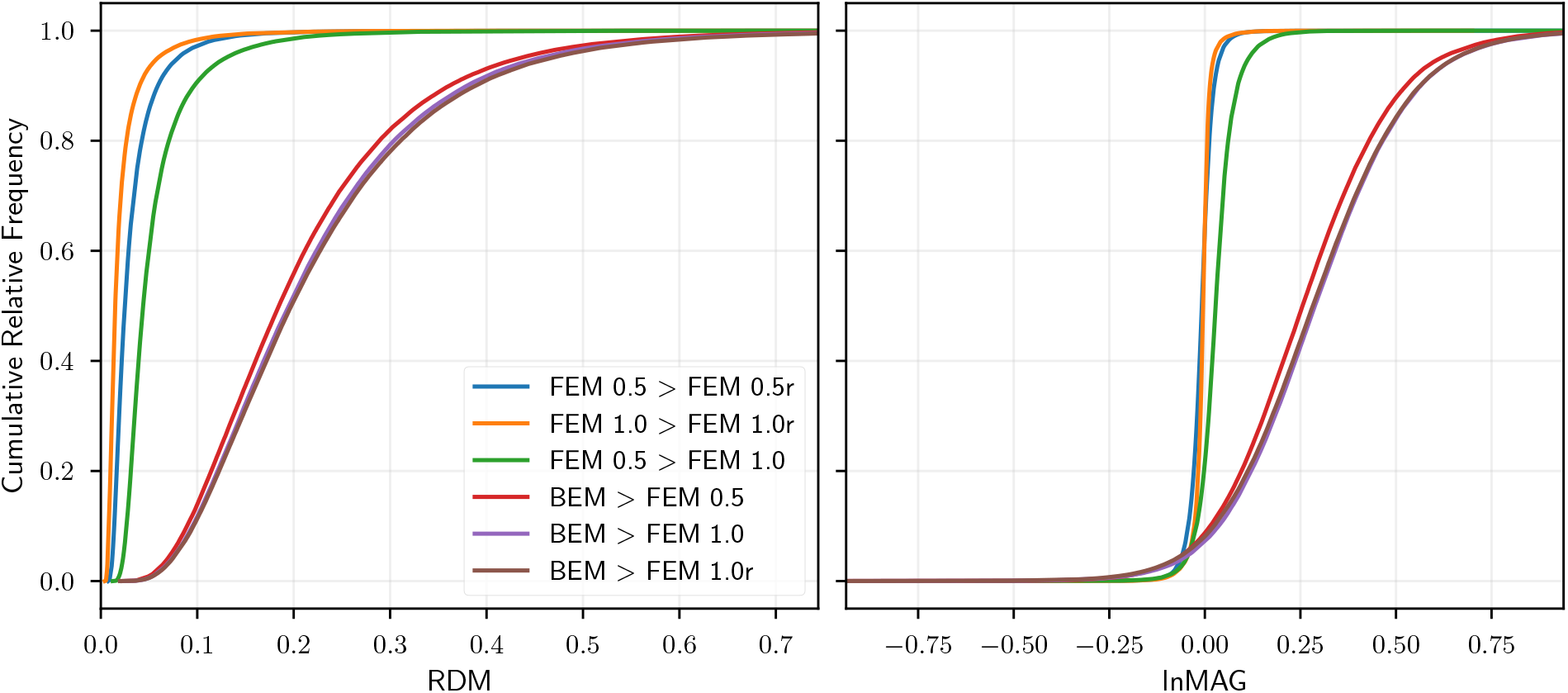
CRF of RDM and lnMAG for FEM comparing five-compartment FEM models of different resolution (0.5 and 1.0 nodes/mm^2^), refined (i.e., upsampled) versions of these models, and a three-layer BEM model. The model on the righthand side of the greater-than sign is the reference.

**Figure 5.**
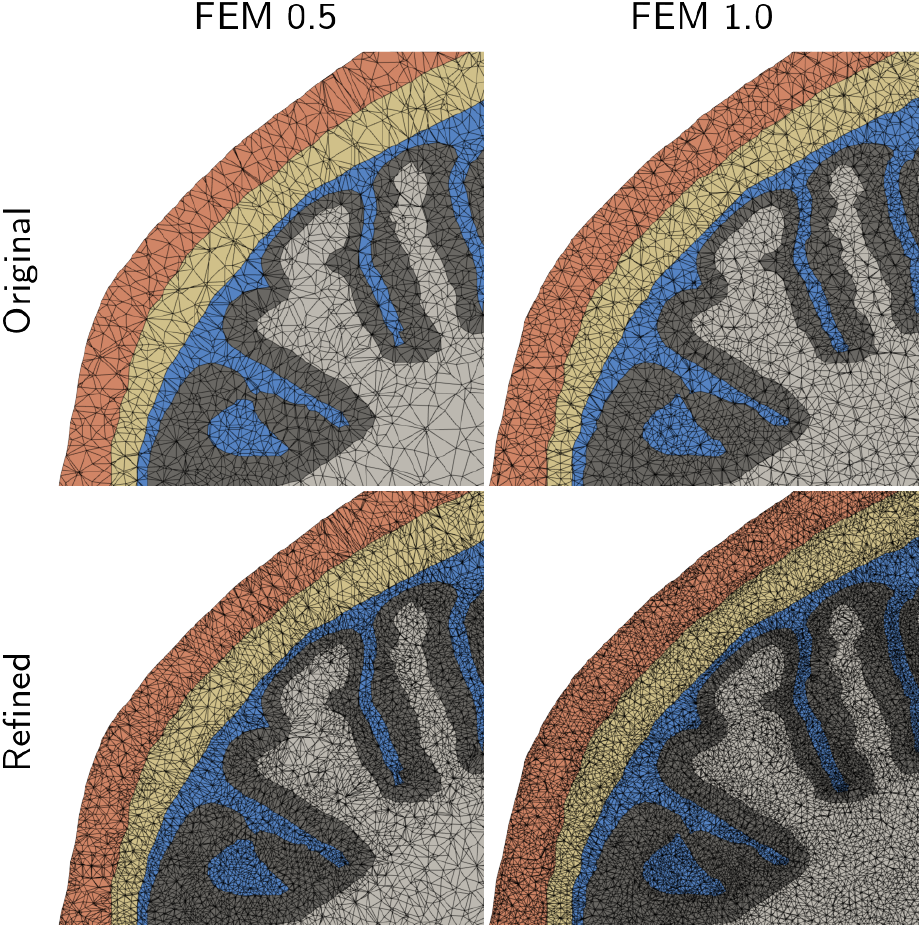
Zoomed view of a coronal slice showing the effect of resolution and refinement on the head models. The corresponding effects on the forward solution can be found in figure 4.

**Figure 6.**
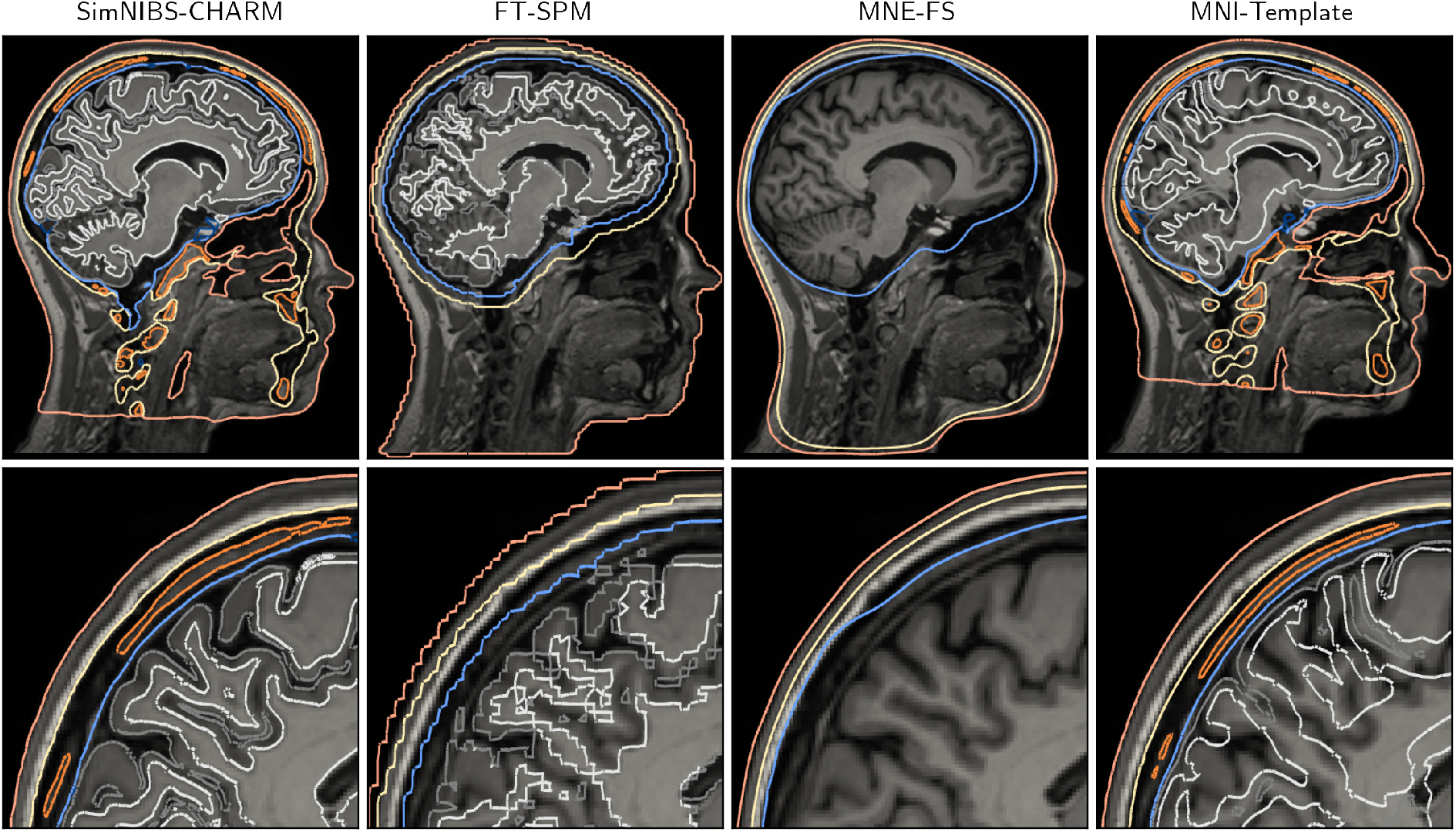
Tissue compartments as estimated by each of the pipelines in the anatomy study for subject one. It is apparant that MNE-FS does not capture the inner skull very well for this subject. The skull thickness varies substantially but this does not seem to agree with the structural MRI scan. Also, the lower part of the head is mostly modeled as skull as the outer skull surface is obtained by eroding the skin surface. Furthermore, the skull thickness in FieldTrip-SPM is almost constant throughout the volume (e.g., thickness is overestimated around the occipital cortex). MNI-Template captures the skull surprisingly well in this subject but the brain tissues are obviously not accurate.

**Figure 7.**
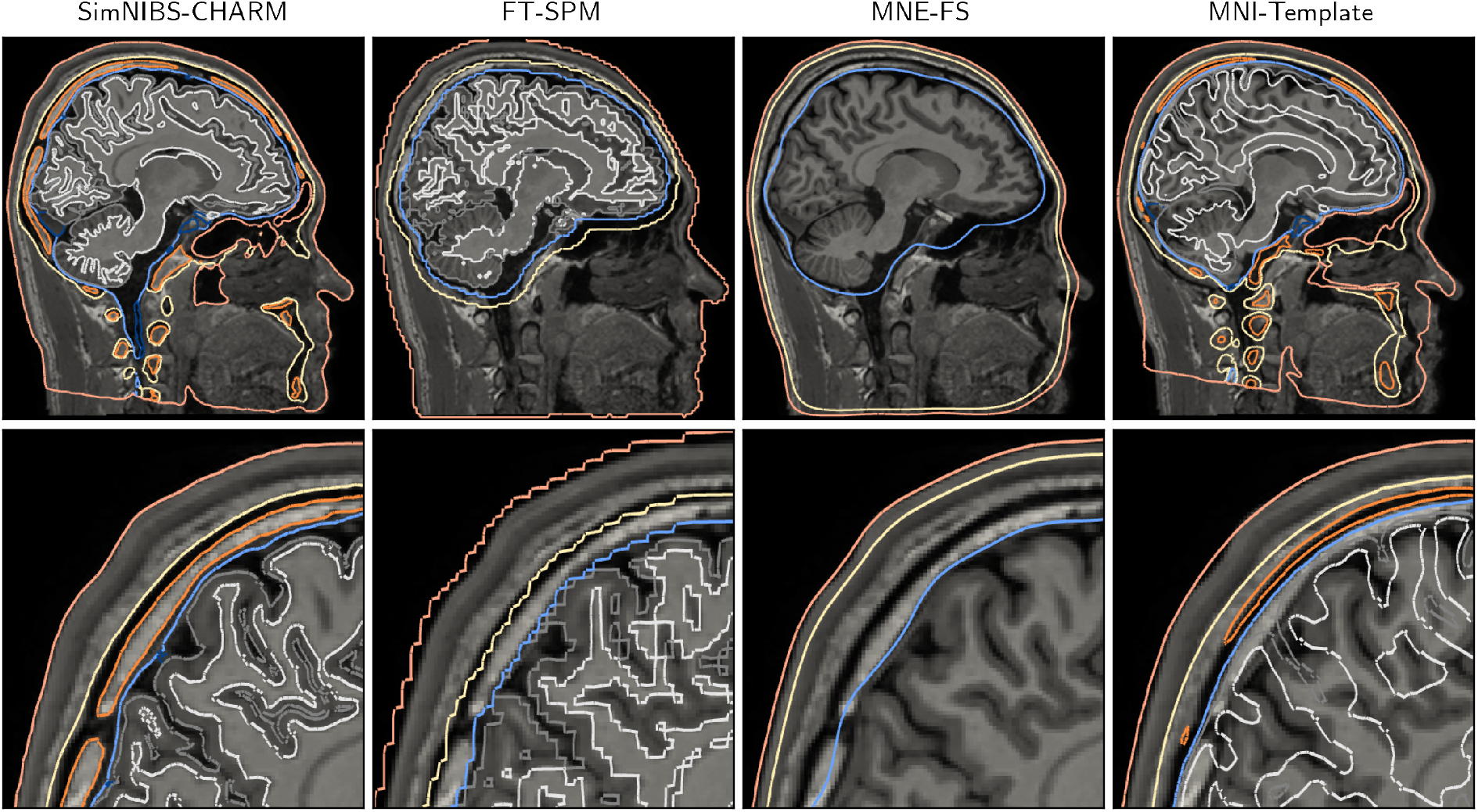
Tissue compartments as estimated by each of the pipelines in the anatomy study for subject two. MNE-FS captures the inner skull better in this subject compared to figure 6, however, the skull thickness is not well modeled. FieldTrip-SPM captures inner skull and thickness well except for the prefrontal part. MNI-Template displaces the skull outwards in the zoomed-in view whereas other parts are modeled more accurately. The spongy bone, however, is not.

### 3.2. Impact of Anatomical Accuracy on the Forward Solution

Figures 6 and 7 show the surfaces extracted using each pipeline overlaid on the T1-weighted image for two subjects. It is not our intention to do a systematic assessment of the anatomical accuracy of the models as this has been done previously (Nielsen et al., 2018; Puonti et al., 2020). However, we believe this to be representative of the differences between the pipelines we have seen in the data.

Generally, we found high anatomical accuracy of SimNIBS-CHARM whereas substantial simplifications of the modeling of the CSF and bone compartments are apparent in FieldTrip-SPM. As a result, FieldTrip-SPM is not particularly accurate for non-brain tissues. MNE-FS collapses white matter, gray matter, and CSF into one brain compartment. Additionally, we found that the BEM surfaces were not very accurate (as evidenced by the need for us to correct intersections between these surfaces). MNI-Template does not capture the individual structure of the brain tissues very well. However, in several cases it actually provided a decent fit of the skull compartment, whereas in others it was not accurate at all. We provide similar figures to figures 6 and 7 for more subjects in section S2 figs. S2.4 to S2.7 in the supplementary material.

Figure 8 shows CRF of the error metrics, RDM and lnMAG, using the forward solution associated with the manual segmentations as reference. In terms of topographical errors, SimNIBS-CHARM shows best performance followed by FieldTrip-SPM, MNI-Template, and finally MNE-FS. As to the magnitude errors, SimNIBS-CHARM and MNI-Template are both centered around zero with the latter having a larger standard deviation than the former. FieldTrip-SPM and MNE-FS generally show decreased and increased sensitivity, respectively. The effect is most severe for MNE-FS both in terms of the mean and standard deviation.

**Figure 8.**
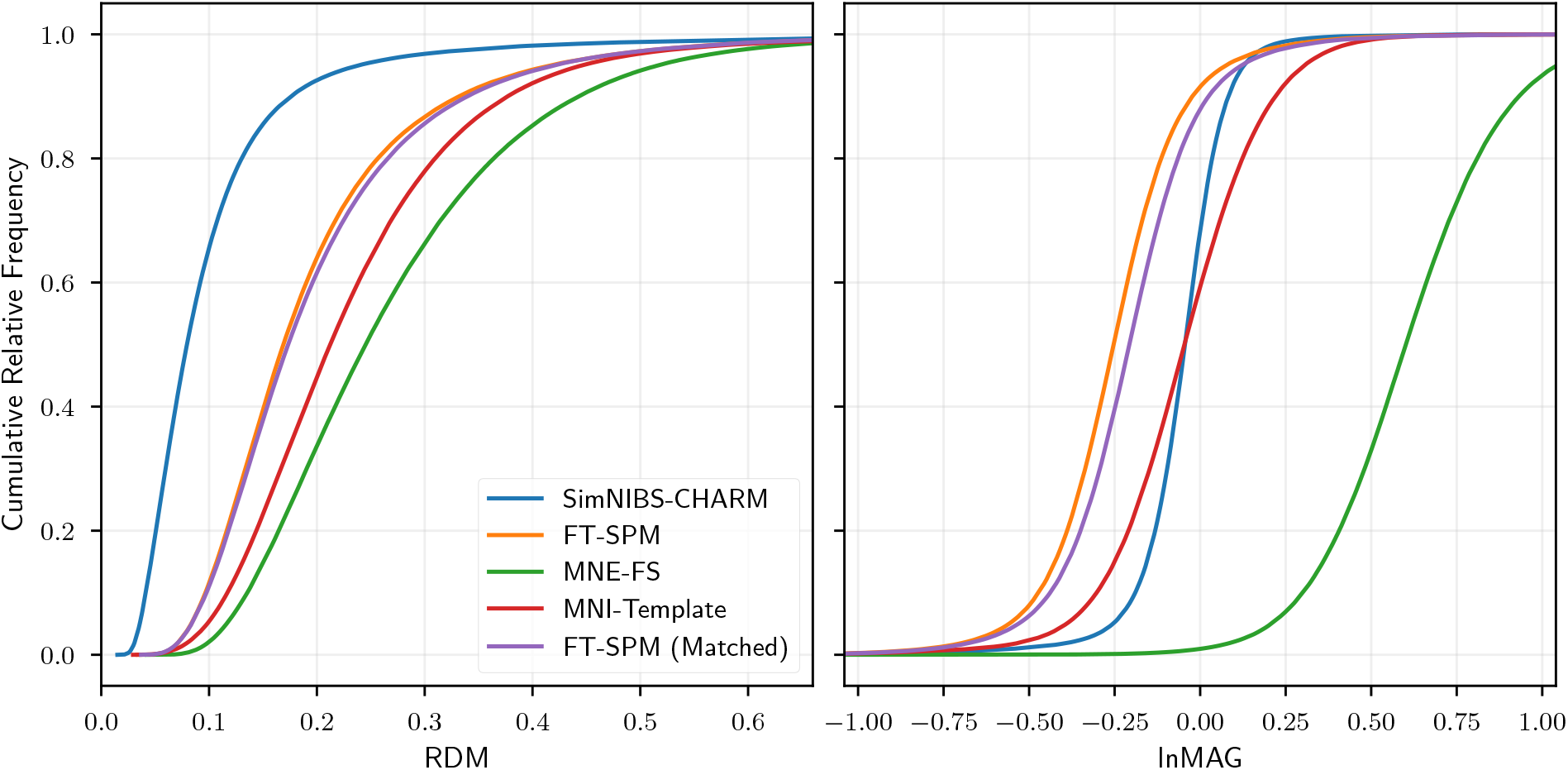
CRF of RDM and lnMAG for each model. FieldTrip-SPM (Matched) is a comparison of the FieldTrip-SPM model with a reference with matched conductivities (see section 4.2.2) allowing us to disentangle effects of geometry and conductivity.

Figures 9 and 10 present the spatial distribution of the mean RDM and lnMAG on the *fsaverage* surface. From figure 9 we see slightly elevated errors in the orbitofrontal area for SimNIBS-CHARM but otherwise low errors. The spatial pattern for FieldTrip-SPM is similar, however, the effects are more pronounced and also extends into the temporal cortex. MNE-FS shows high errors on the gyral crowns both for superior and deep sources. For MNI-Template we see mostly errors comparable to those of FieldTrip-SPM, but with a few hot spots and also slightly increased errors in parts of the parietal cortex. From figure 10 we that SimNIBS-CHARM generally has the lowest overall errors. FieldTrip-SPM shows decreased sensitivity to deep, occipital, and temporal sources whereas MNE-FS shows increased sensitivity everywhere, particularly for shallow sources on the gyral crowns.

**Figure 9.**
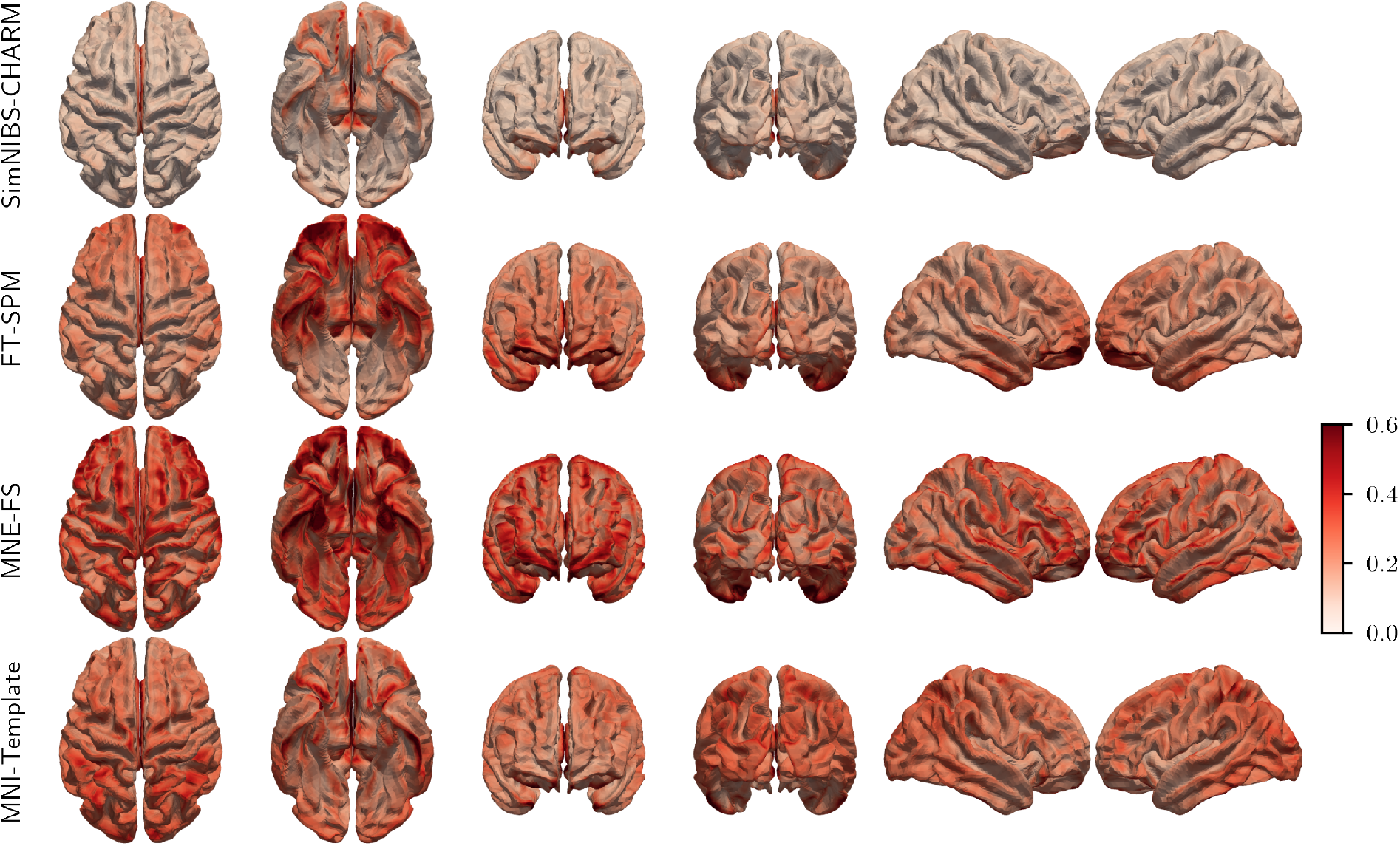
Spatial distribution of mean RDM for each model.

**Figure 10.**
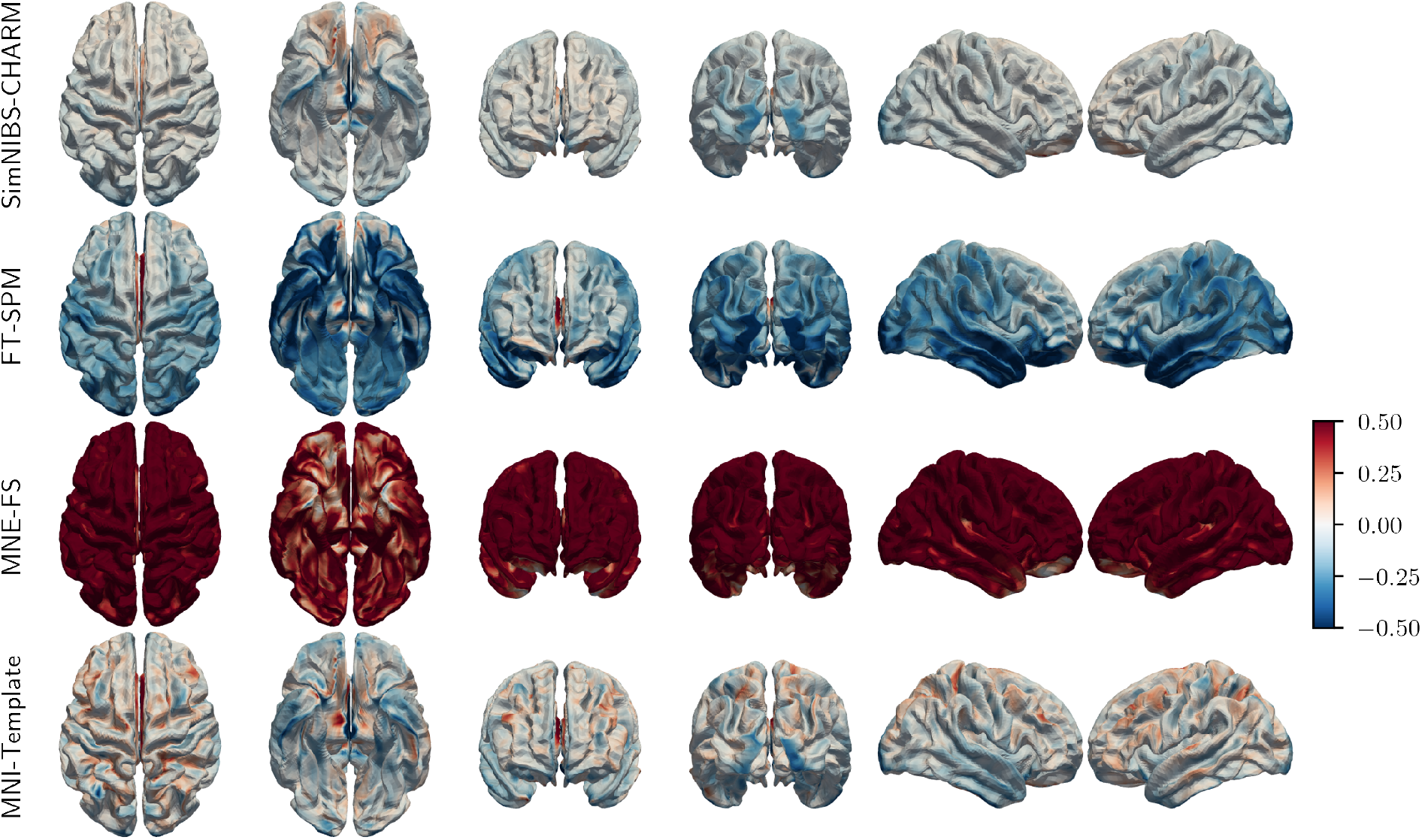
Spatial distribution of mean lnMAG for each model.

We expect the magnitudes to be sensitive to the ohmic tissue conductivities. Since we used the default values in each pipeline—and because the conductivities are the same in SimNIBS-CHARM and MNI-Template as in the reference model—it is not surprising that these are similar in terms of sensitivity. On the other hand, differences in conductivity is a potential confounder for the comparison of the FieldTrip-SPM and MNE-FS models to the reference. In an attempt to disentangle such effects, we created another reference model with conductivities similar to those of FieldTrip-SPM^11^ and computed RDM and lnMAG. From figure 8, we see that most of the differences remain even after this correction, and so we believe that our statements above are indeed valid. Due to the vast geometrical simplifications in the MNE-FS model (collapsing brain tissue and CSF into one compartment) and their known effects on the forward solution (Vorwerk et al., 2014), we did not do a similar analysis with the MNE-FS model as we feel the geometrical adjustments to the reference model would be too large.

### 3.3. Impact of Electrode Positions on the Forward Solution

We first report errors on the channel positions before describing the resulting effects on the forward solution.

Figure 11 is a density plot of the Euclidean distance between electrode positions estimated using the different methods and the digitized positions across all subjects. It is evident that Man-Template generally performs worse than Custom-Template and also that the particular transformation method of Custom-Template does not have a strong influence on the accuracy of the electrode positions.

**Figure 11.**
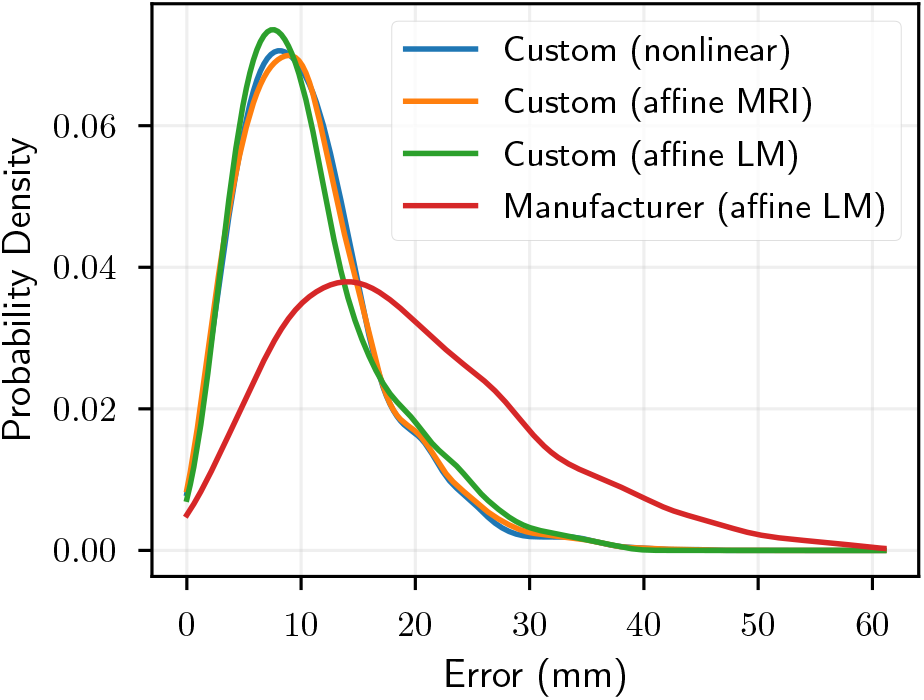
Density of errors in electrode position as measured by the Euclidean distance between each model and the digitized positions. The parenthesis denotes the registration method where “LM” means that the registration was based on landmarks.

Figure 12 shows the spatial distribution of the mean error over channels. We see that the errors in the posterior areas (mostly occipital but also parietal and temporal) are large for Man-Template whereas this is much less severe for the Custom-Template. Likewise, the standard deviation is also larger for Man-Template, again in posterior but also anterior areas. Interestingly, although the mean error is similar across the different transformation methods for Custom-Template, the standard deviation is not. The MRI-based coregistrations show larger variability in the anterior parts of the head (suggesting that other parts of the image are driving the registration). The landmark-based registration exhibits larger variability in the posterior regions (perhaps unsurprisingly given that the landmarks used here are the nasion, LPA, and RPA).

**Figure 12.**
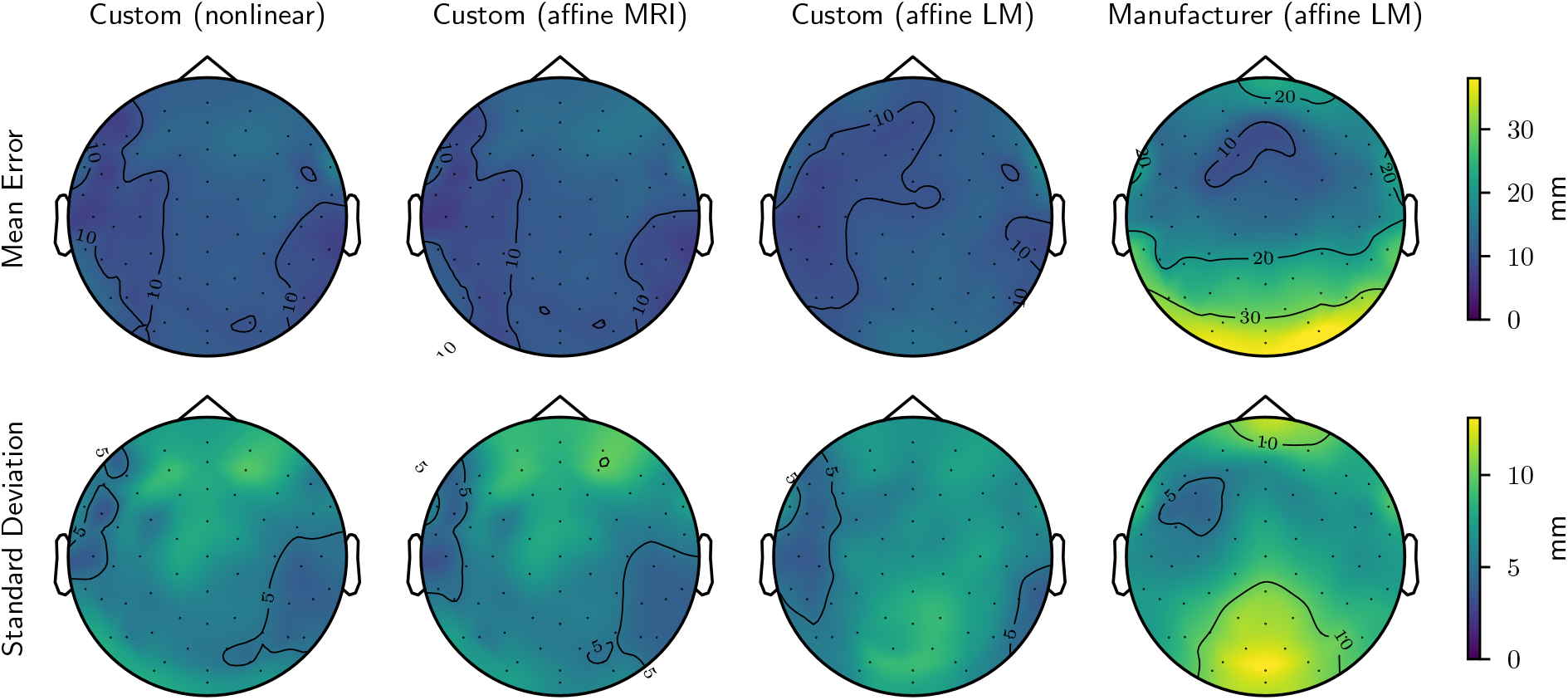
Spatial distribution of errors (mean and standard deviation) in electrode position as measured by the Euclidean distance between each model and the digitized positions. The parenthesis denotes the registration method where “LM” means that the registration was based on landmarks.

To determine potential spatial bias in the template registrations, we show the absolute errors along each axis in figure figure 13. In general, it seems that errors in anterior and posterior areas are largest in the *z* direction and errors in the superior areas are largest in the *y* direction. This suggests that the errors here are mostly due to an anterior-posterior (AP) shift (i.e., a rotation around the *x*-axis). Interestingly, errors in the most inferior electrodes in the temporal areas are primarily due to shifts along the AP direction and not up or down. Finally, please note the different scales of Custom-Template and Man-Template and that errors may be exaggerated close to the edges of the head contour due to extrapolation.

**Figure 13.**
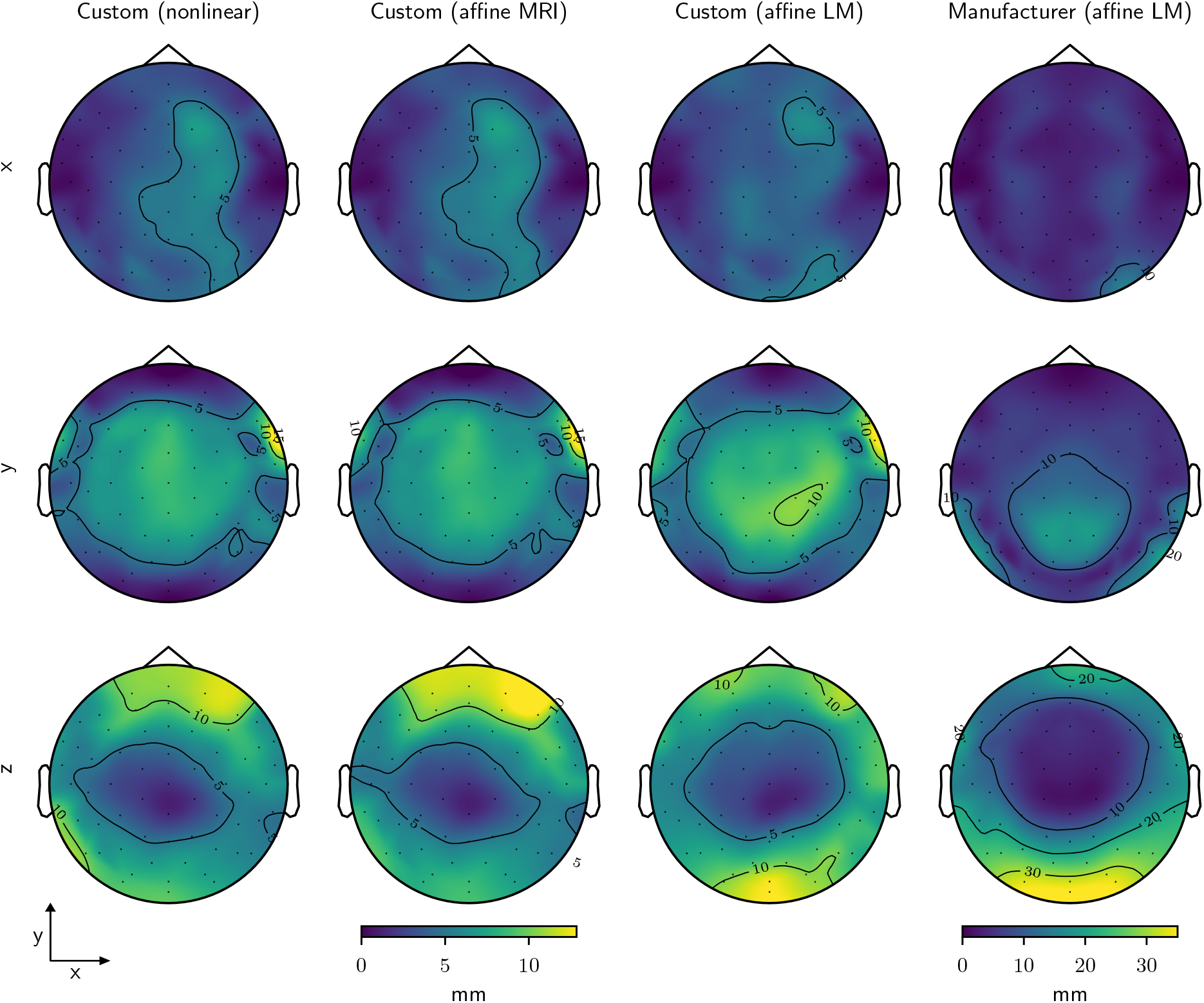
Spatial distribution of mean errors in electrode position as measured by the absolute difference between each model and the digitized positions. Please note the different scale of the rightmost column. The parenthesis denotes the registration method where “LM” means that the registration was based on landmarks. Positive *z* direction is out the page.

For both Custom-Template and Man-Template, we found that channel errors in occipital and frontal regions were mostly along the *z*-axis and along the *y*-axis for superior areas. To investigate the extent to which errors were due to bias (i.e., a systematic difference between the template and the average positions across subjects) or variability in the reference positions (i.e., the standard deviation of each position, e.g., due to inter-individual differences in cap placement), we transformed the reference positions for all subjects to standard space and computed bias and standard deviation for each position. The standard deviations (figure 14) showed the same pattern as that observed in figure 13, suggesting more variability in AP alignment of the cap compared to aligning it laterally. On the other hand, we also found a higher bias at some positions than others (figure 15). For example, the digitized positions of the electrodes between the ears and the eyes were, generally speaking, more anterior compared to Custom-Template.

**Figure 14.**
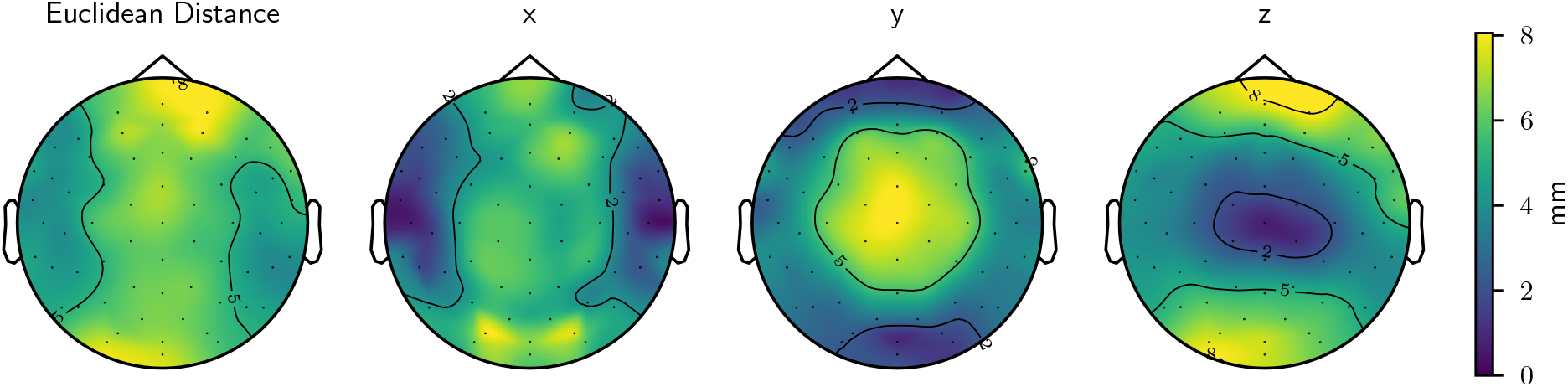
Standard deviations of the digitized positions in the electrode study calculated using the overall Euclidean distance or distances along each major axis. This illustrates the variability of the reference positions.

**Figure 15.**
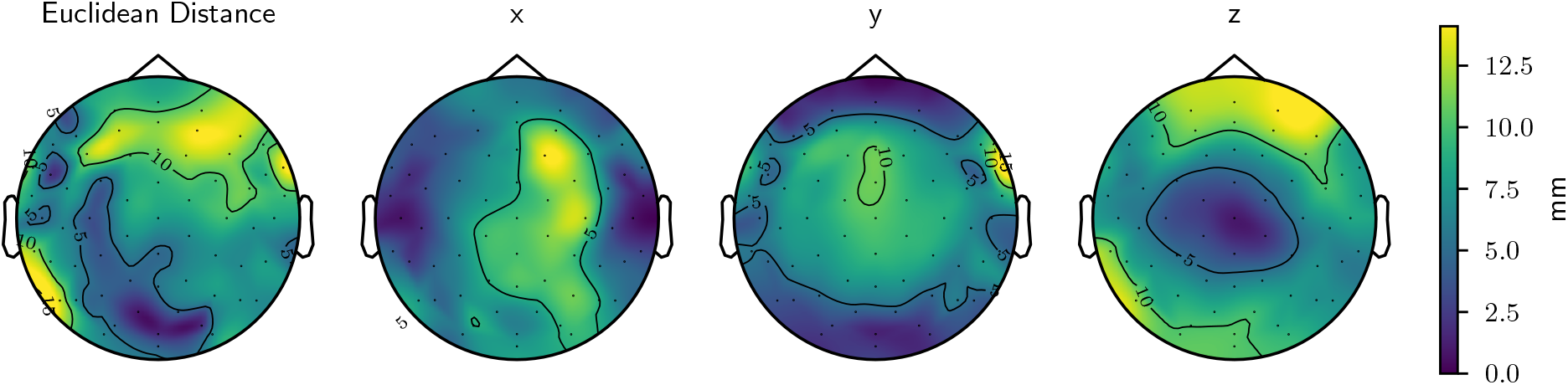
Distance between the average of the digitized positions in the electrode study and the positions of the custom template calculated using the overall Euclidean distance or distances along each major axis. This illustrates the bias of the template compared to the reference positions.

We also show all digitized electrode positions in standard space on the MNI surface template (figure 16) from which it is apparent that the positions towards the back are lower in the left side compared to the right side. This points to some kind of misalignment, perhaps due to the way the chin strap was fastened being different between subjects and the MNI template head. As such, it is not clear whether the observed bias is due to suboptimal (biased) cap placement in the data set used here or because the template fails to capture how a cap fits on an actual human head.

**Figure 16.**
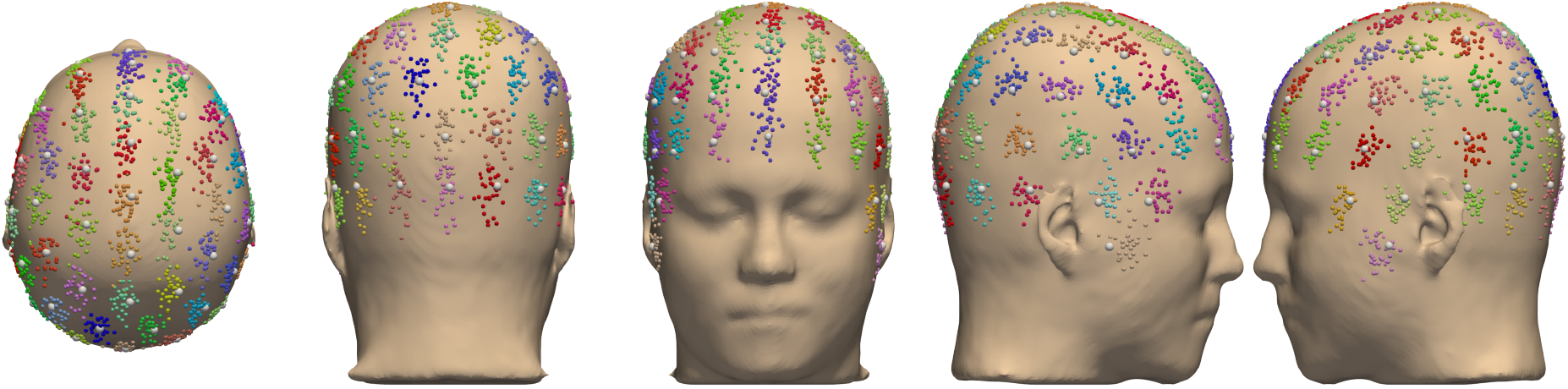
Our custom template positions (white spheres) and the digitized positions for each subject (colored spheres) in MNI coordinates.

Figure 17 shows CRF of the error metrics, RDM and lnMAG, for the forward solutions using the solution associated with the digitized positions as reference. It is evident that the RDM errors are generally higher for Man-Template than for Custom-Template whereas the errors for MNI-Digitized fall between the two. As to lnMAG, Custom-Template is again better than Man-Template but here MNI-Digitized is clearly worse than the other two as evidenced by the heavy tails.

**Figure 17.**
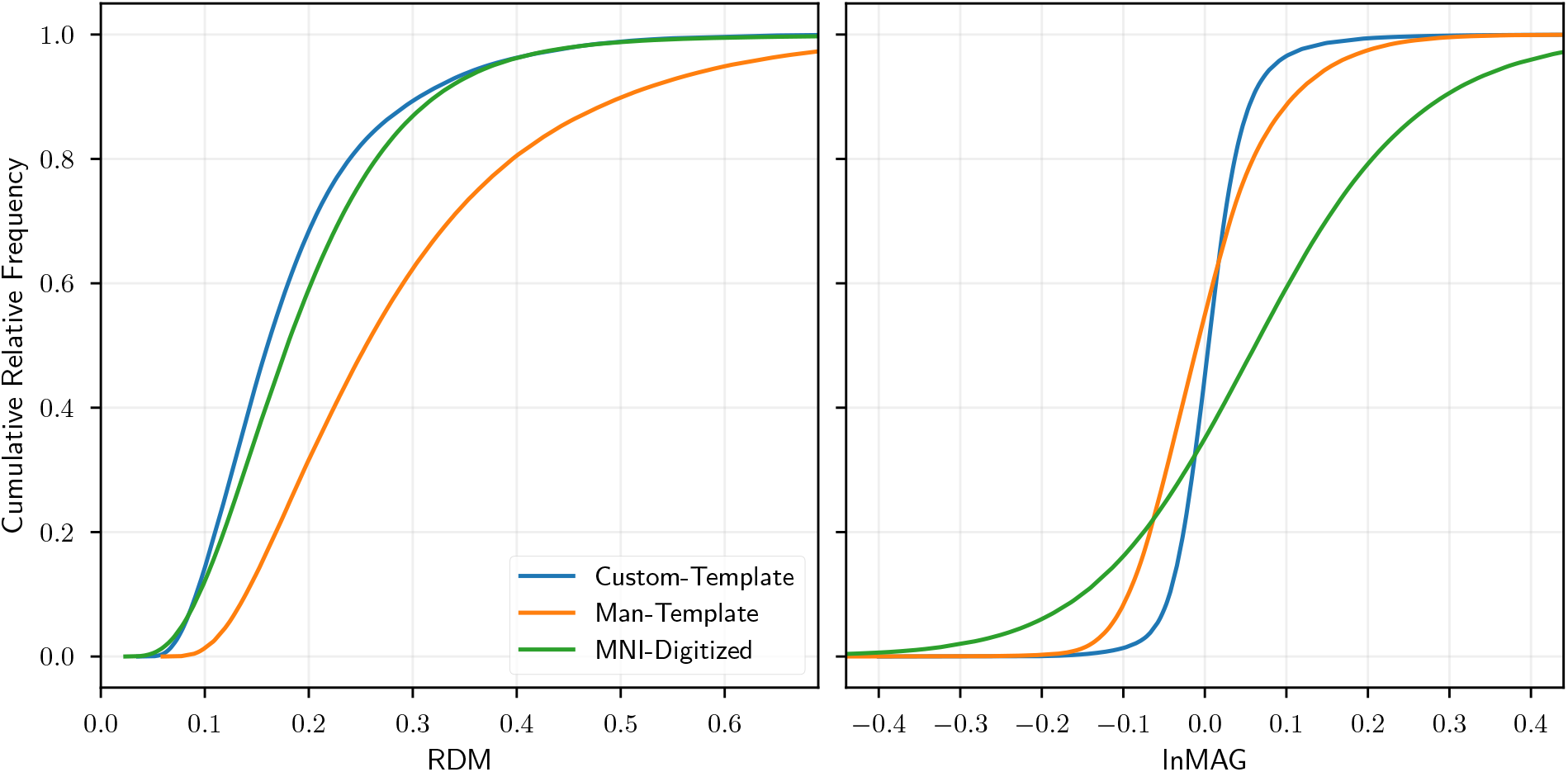
CRF of RDM and lnMAG for each model.

Figures 18 and 19 present the spatial distribution of the mean RDM and lnMAG on the *fsaverage* surface. From figure 18, we see slightly elevated topographical errors for Custom-Template in orbitofrontal and occipital errors. However, this is more pronounced in Man-Template which also shows increased errors in parietal and temporal regions. The errors of MNI-Digitized are generally similar to those of Custom-Template. However, we see several hot spots of increased error, often on the gyral crowns. From figure 19, we see slightly increased magnitudes for Custom-Template in frontal and right temporal regions. Generally, Man-Template shows increased and decreased magnitudes for deep and shallow sources, respectively. Again the magnitudes are elevated in the right temporal region. MNI-Digitized shows large magnitude errors with increased and decreased sensitivity to sources on the gyral crowns and sulcal walls, respectively. Decreased sensitivity is also observed for orbitofrontal and occipital sources.

**Figure 18.**
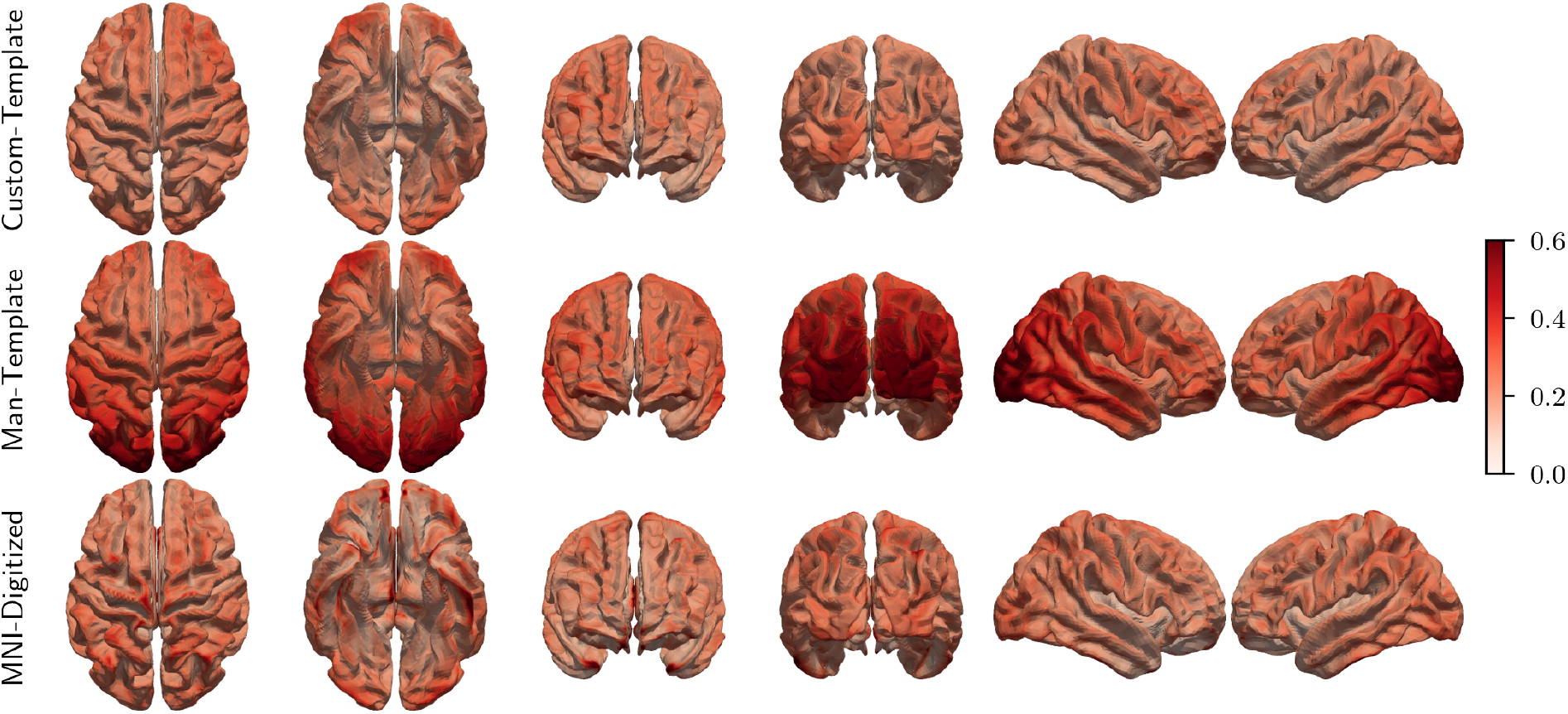
Spatial distribution of mean RDM for each model.

**Figure 19.**
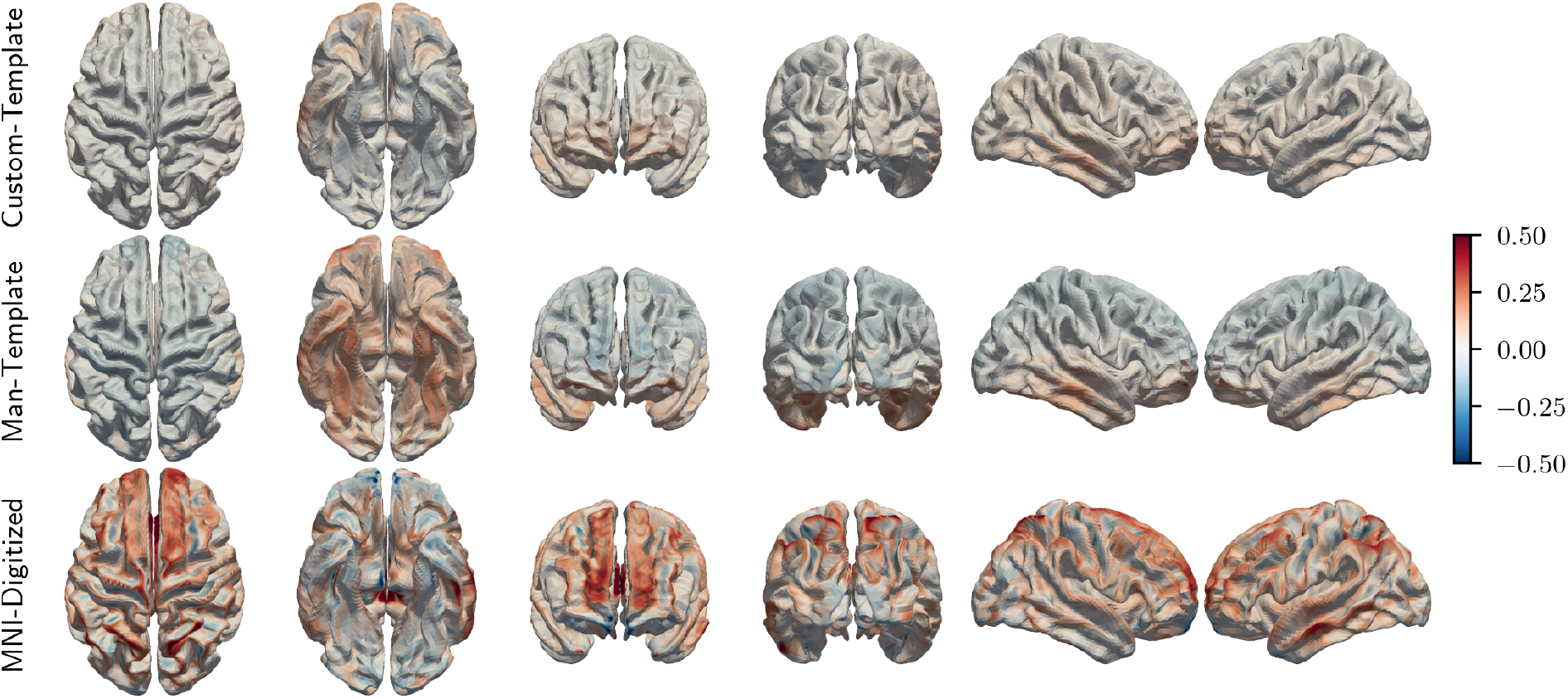
Spatial distribution of mean lnMAG for each model.

## 4. Discussion

### 4.1. Validation of SimNIBS for EEG

The simulations in spherical models, assuming sufficient model resolution of about 0.5 or 1.0 nodes/mm^2^, showed that the numerical errors in topography and magnitude are low and similar to those of existing FEM implementations for EEG (Vorwerk et al., 2012). The same was true for simulations using a realistic geometrical models. In addition, by comparing numerical errors of our FEM to errors incurred by not distinguishing different tissue compartments in the BEM head models (see figures 3 and 4), we show that the latter are about ten times those of the former. Hence, we suggest that in general the decrease in numerical accuracy associated with FEM is offset by its flexibility when it comes to modeling complex anatomical structures.

Close inspection of figure 2 reveals that there is a small bias in the magnitude in the SimNIBS simulations. Decreasing the thickness of the electrodes (thus minimizing potential drop over the electrode itself) and matching the conductivity to that of skin (since high conducting electrodes effectively makes the skin compartment a better conductor) removed this effect. Thus, we attribute this to the electrodes being meshed in SimNIBS whereas the analytical solution assumes point sensors. One could argue that the correct approach is to model the electrodes. However, the effects are very small in practice.

As to the source model used in SimNIBS, calculating the electric field in each element is similar to the partial integration approach (Vorwerk, 2016) with dipoles along each axis. However, to improve the accuracy of the estimate, we use a postprocessing procedure (SPR) which consists in fitting a smooth function such that the value of a node is given by a linear combination of its neighboring elements. Although different from the St. Venant approach (Vorwerk, 2016), we found the numerical accuracy to be similar.

### 4.2. Impact of Anatomical Accuracy on the Forward Solution

#### 4.2.1. Topographical Errors

Interpreting the topographical errors reflected by RDM can be difficult since the quality of the anatomical models may vary from subject to subject. However, we will summarize some general observations about the different models.

Both FieldTrip-SPM and MNE-FS showed elevated topographical errors in orbitofrontal and deep temporal areas. Given the close proximity of such sources to the facial area and the fact that neither of these models attempt to characterize the complex anatomy in this region, we suspect that such simplifications may explain the increased RDM values here. Since the anatomy is simpler around the occipital cortex, the modeling decision of FieldTrip-SPM to simply enclose the brain with a certain amount of bone (and CSF) seems more appropriate here. The same holds for MNE-FS which also showed large RDM values for superficial sources in the superior part of the brain. Similar observations were made by Vorwerk et al. (2014) when distinguishing CSF and brain tissues. The inaccuracy of the inner and outer skull surfaces in MNE-FS likely exacerbates this issue (Lanfer et al., 2012).

In line with the better anatomical accuracy, the topographical errors of SimNIBS-CHARM are generally small compared to the other models (see figures 6 and 7). MNI-Template seems slightly better than MNE-FS but worse than the others. This seems to highlight the importance of accurately modeling CSF since the brain tissue compartments in MNI-Template are generally very different from the individual anatomy.

#### 4.2.2. Magnitude Errors

As with the topographical errors, the magnitude errors of SimNIBS-CHARM are low whereas they are slightly higher for MNI-Template (heavier tails in the CRF curve). The errors of MNE-FS and FieldTrip-SPM are larger still and biased towards higher and lower sensitivity, respectively.

The magnitude errors in FieldTrip-SPM suggest that sensitivity is generally decreased, particularly for deep sources. From figures 6 and 7 it is evident that the models tend to contain too much CSF, particularly in the deep and temporal areas where bone is also too thick, which will increase shunting effects. MNE-FS shows the opposite effect, i.e., an overall increase in magnitude, most prominently for sources close to the sensors. MNE-FS does not model the CSF compartment explicitly but rather collapses white matter, gray matter, and CSF into a single compartment with an adjusted conductivity. Thus, the conductivity gradient between the brain and bone compartments is smaller and hence we expect less current shunting. These results are in line with Vorwerk et al. (2014) in terms of the relevance of distinguishing CSF from brain tissue as well as white matter from gray matter.

### 4.3. Impact of Electrode Positions on the Forward Solution

Similar to Homölle and Oostenveld (2019), we also found that channel errors were generally much larger for Man-Template than Custom-Template in occipital and parietal areas suggesting that approximating the head shape with a sphere is problematic when the sphere is aligned using frontal and temporal landmarks. Including the inion in the coregistration could potentially help, although we did not test this. On average, we did not find similar effects in posterior regions when transforming Custom-Template using landmarks (although standard deviations were higher), suggesting that the misalignment of Man-Template is in fact due to this layout being a poor fit of the actual positions in these areas.

We also investigated the extent to which the errors observed in Custom-Template were due to bias (i.e., systematic differences to the reference positions) or variability (in the reference positions). We found that the errors were partly due to a bias of the template but also that some electrodes were less consistently placed on the individual subjects. We did not do a similar analysis for Man-Template.

Given this bias and variability in the electrode positions, it is not reasonable to expect a perfect fit of the template to all subjects. We noticed that the fit was particularly bad in a few cases where the cap was clearly not aligned well (e.g., rotated around the *x* or *z* axis).

#### 4.3.1. Topographical errors

In line with the mean error observed on the channel level, topographical errors in the forward solutions, as estimated by RDM were more severe for Man-Template than Custom-Template. For the former, errors were high in occipital and posterior areas. Errors were higher on the gyral crowns (i.e., where the sensitivity is high) than the sulcal walls and valleys, something which is most clearly seen for Man-Template.

Topographical errors for MNI-Digitized are slightly higher than Custom-Template but smaller than Man-Template suggesting that electrode positions are important in determining such errors and that template electrode positions as supplied by the manufacturer result in higher RDM values compared to a template anatomy with digitized positions. This is in line with the results of Homölle and Oostenveld (2019).

We also see a few RDM hot spots in MNI-Digitized which, as they stand out on a group level, are likely due to local inaccuracies in this model. We were not able to identify any obvious errors upon visual inspection of the final segmentation and mesh though.

#### 4.3.2. Magnitude errors

We observed that Man-Template showed increased and decreased magnitudes for deep and superior sources, respectively. Visual inspection of the final positions showed that electrodes were generally placed further down on the head model, thus giving increased sensitivity to deep (or ventral) sources, and, conversely, decreased sensitivity to superior sources (most notably parietal, occipital, and prefrontal) as the montage was spread more thinly across the head. Increased sensitivity to temporal sources was also observed as these are generally not very well covered by electrodes.

Magnitude errors were much larger for MNI-Digitized. In particular, we observed increased sensitivity to sources on the gyral crowns and decreased sensitivity to sources on the sulcal walls. Since the MNI152 T1-weighted template represents a standard anatomy, it is generally quite blurred. As a result, fine details such as gray matter curvature and sulcal structure are lost. Visual inspection (see figure 20) showed that the gray matter compartment of the final head model was much smoother than for individual subjects, hence sources on sulcal walls and in sulcal valleys were surrounded almost exclusively by gray matter and not highly conducting CSF as was the case in the models based on the individual anatomy. Additionally, we also found that the CSF layer was much thinner compared to the individualized models resulting in less current shunting, particularly for sources on the gyral crowns to which sensitivity was increased (see figure 20).

**Figure 20.**
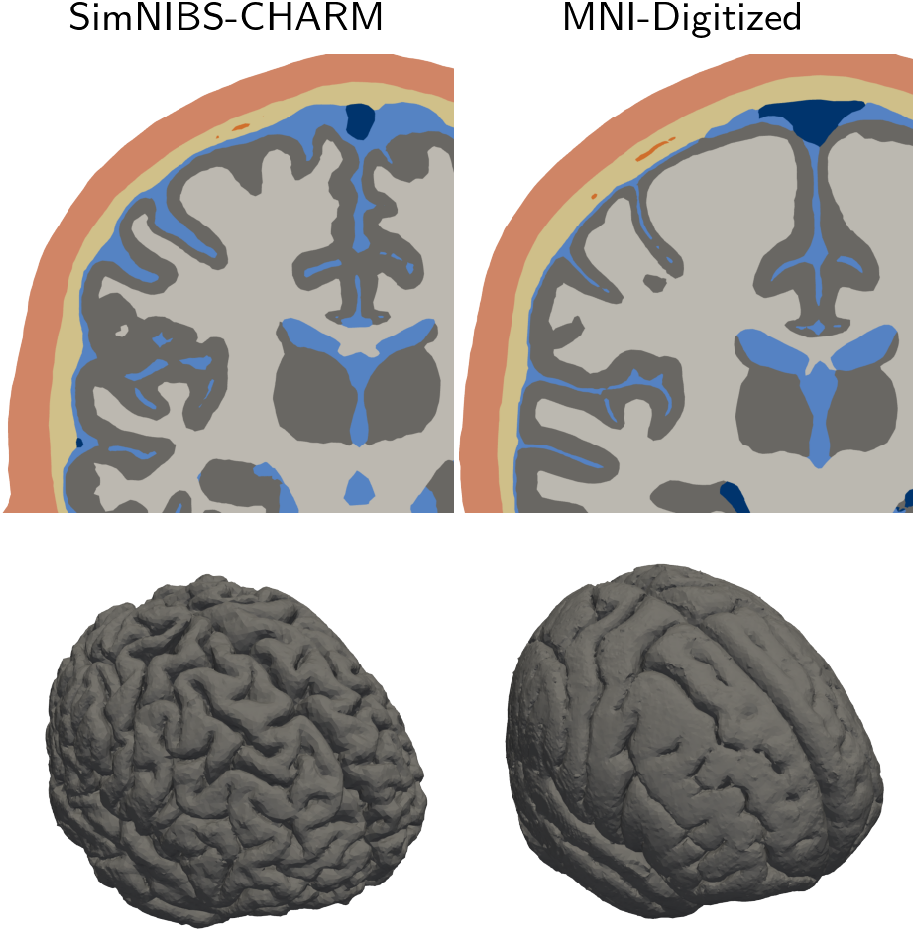
The top row illustrates the reduced amount of CSF between the cortex and the inner skull in MNI-Digitized compared to the SimNIBS-CHARM for a particular subject (the MNI-Digitized model is almost identical for all subjects except for the nonlinear deformation). The bottom row shows the corresponding gray matter compartments.

### 4.4. Effect on Inverse Solution

Despite clear differences in the forward solutions obtained from the different pipelines investigated in this study, the impact on the inverse solution still needs to be established. There is, however, already a body of literature suggesting that the forward model is indeed an important factor that influences source reconstructions. Below, we discuss this in relation to the effects observed in the current study.

As mentioned in section 1, the modeling of the skull compartment has received much attention in the EEG community and from figures 6 and 7 it is apparent that there are clear differences in how the pipelines model this. The thickness is not accurately captured in FieldTrip-SPM and MNE-FS whereas SimNIBS-CHARM is able to delineate the skull with decent accuracy (Puonti et al., 2020). However, previous studies report only minor source localization errors (less than 5 mm) due to such erroneous thickness estimation (Chauveau et al., 2004; Lanfer et al., 2012). It is also apparent that FieldTrip-SPM and MNE-FS simplifies the modeling of the inferior part of the skull quite substantially. This was shown by Lanfer et al. (2012) to result in large errors (more than 1 cm) for sources close to the base of the skull. A related point has been made by studies showing that failure to model skull openings (foramina) (Fiederer et al., 2016) and sutures (McCann and Beltrachini, 2022) may decrease localization accuracy particularly for sources in the vicinity of such errors. Finally, Montes-Restrepo et al. (2014) showed that low quality skull segmentations (generated from T1-weighted images compared to CT scans) resulted in increased source localization errors of up to 1 cm, but only for sufficiently low noise levels.

The human skull, however, is not a homogeneous compartment since it is comprised of both spongy (soft) and compact (hard) bone. SimNIBS-CHARM distinguishes between these, which is not the case for FieldTrip-SPM and MNE-FS. Dannhauer et al. (2011) suggests that modeling such inhomogeneity is important and that failing to do so may result in increased localization errors up to 1 cm. On the other hand, using anisotropic modeling of the skull (different conductivities in the radial and tangential directions) to account for the layered structure of the bone may not lead to any noticeable improvements and may even deteriorate performance in areas of complex geometry (Dannhauer et al., 2011; Montes-Restrepo et al., 2014). We would like to point out though—irrespective of whether or not the particular distinction is important for source localization *per se—*that within the Bayesian segmentation framework used by CHARM, modeling compact and spongy bone separately may be beneficial for the overall segmentation result because spongy bone and skin often look similar on a T1-weighted image. Thus, this might improve the segmentation quality in general, particularly in subjects where spongy bone is very prominent or where the image quality is low (e.g., corrupted by fat shift) (Nielsen et al., 2018; Puonti et al., 2020).

Another important aspect of forward modeling is whether or not to distinguish between different brain tissues and CSF or simply model it as one compartment. The latter strategy is employed by MNE-FS (and most other pipelines using BEM) whereas both FieldTrip-SPM and SimNIBS-CHARM includes white matter, gray matter and CSF. Ramon et al. (2006) concluded that the CSF compartment is important for shaping the observed potential distribution on the scalp and neglecting this increases average source localization errors by a few millimeters. Likewise, Conte and Richards (2021) found that including CSF was the most important addition to a model consisting initially of brain, skull, and scalp and that when doing so, it performed similarly to more complex models distinguishing even more tissue types. The latter suggests that the increased complexity of the SimNIBS-CHARM model (including for example muscle tissue and eyes) may not matter much in terms of source localization accuracy. Similarly, differentiating between white and gray matter was found to be less important compared to CSF (Ramon et al., 2006).

In terms of anatomical accuracy, there is substantial variation as to how well the template model fits each subject. Generally, the brain tissues and CSF do not match at all. However, the inner and outer skull borders are a more reasonable fit. Acar and Makeig (2013) observed median dipole localization errors of about 5 mm when using a realistic four compartment template model warped to the head shape of each subject (as was also done in this study). They found high errors towards the base of the skull since their template was cut just below the nasal area. However, the template used here has similar coverage to that of the individualized models so we expect this to be less of an issue (Lanfer et al., 2012).

In the study investigating the impact of the accuracy of the electrode positions, we found spatially correlated errors concentrated around occipital and parietal areas. This was true for both Custom-Template and Man-Template, but was more prominent for the latter. Similar results were reported by Homölle and Oostenveld (2019), who found average dipole localization errors of 8.5 and 11 mm for custom and manufacturer templates, respectively. Again, particularly high errors for the latter occurred over the above mentioned areas. In an attempt to simulate coregistration errors, Acar and Makeig (2013) tilted electrode positions 5 degrees and found mean localization errors about 6 mm along the direction in which the tilt was performed (e.g., left-right tilt caused larger errors in temporal areas). Thus, spatially correlated errors in electrode locations seem to propagate to source localization errors. On the other hand, Wang and Gotman (2001) found only minimal effect of electrode position on source localization accuracy. however, in this study the simulated errors were uncorrelated between neighboring sensors.

Studies have also suggested that the choice of digitization method may impact inverse solutions. For example, Dalal et al. (2014) reported decreased beamformer sensitivity and slightly increased localization errors when electrode positions were recorded using a suboptimal digitization procedure and Shirazi and Huang (2019) found that digitization errors of about 1 cm could potentially result in source reconstruction errors of up to 2 cm. In the latter study, they report degraded localization accuracy using a template of electrode positions. However, it is not clear from the study exactly how these positions were obtained, and, as suggested by our results and those of Homölle and Oostenveld (2019), this may significantly impact the accuracy.

While the literature reviewed here suggests that errors in the head model and electrode positions used for forward modeling affect the accuracy of the inverse solution, this influence may be modulated by other factors such as the SNR of the measurements and the inverse modeling approach used. For example, Montes-Restrepo et al. (2014) show that source localization accuracy benefits from accurate skull modeling based on CT, however, they also found that differences between CT and MRI based skull models disappeared as noise levels increased (e.g., going from SNR of 10 dB to SNR < 5 dB). Ramon et al. (2006) investigated the effect of distinguishing different tissue compartments (e.g., white matter from gray matter or merging CSF, white matter, and gray matter) on source localization accuracy. They found that differences between models only started to emerge above an SNR of approximately 5 dB. Thus, a sufficient SNR seems to be required to benefit from more complex forward modeling. Both of these studies used dipole fitting for localizing sources. On the other hand, distributed inverse methods are generally expected to be more robust to forward modeling errors at the expense of being less focal (even in ideal conditions) as well as suffering from depth bias (Stenroos and Hauk, 2013).

### 4.5. Limitations

The comparison of different pipelines is likely biased in favor of SimNIBS-CHARM since the tissue priors used by the CHARM pipeline was built from this dataset. Although we used alternative priors from a four-fold split such that a particular subject never contributed to the prior used for generating its own segmentation, we might still expect some bias. On the other hand, it is clear from simple visual inspection of the segmentations that SimNIBS-CHARM is more accurate than FieldTrip-SPM and MNE-FS. As such, we believe the observed differences are robust and will generalize to other MRI datasets as well. Similar to SimNIBS-CHARM, the results of the model based on a template anatomy might also be biased given that it contains the same tissue classes and was constructed from the same pipeline. To minimize bias, we again used priors from a four-fold split.

The reference models used in the anatomy study, although based on manual segmentations, were still not perfect and certain simplifications were made. For example, we failed to model brain conductivity anisotropy. However, assuming that the effect is similar across different pipelines, we do not expect it to strongly affect relative differences. Besides, white matter anisotropy seems to mostly affect certain deep sources and effects are generally small compared to other modeling aspects (Vorwerk et al., 2014).

In the electrode study, we used a dataset collected at our department as part of a previous project. Since electrode positions were digitized in this study, the experimenter may have been less careful with cap placement than if it was known beforehand that a template was to be used to model the electrode positions. As such, they might not constitute the best possible case when evaluating the fit of an electrode template. For example, visual inspection (as well as the standard deviation some of the electrodes across the dataset) suggested that in some subjects the cap was clearly misaligned in the anterior-posterior direction or twisted left or right. On the other hand, we believe that this is likely to happen in an actual experimental setting and as such our results may give a realistic impression of the errors that might be incurred using such methods.

## 5. Conclusion

In summary, it seems that topographical errors, as measured by RDM, are sensitive to errors in the modeled anatomy and the sensor positions, whereas magnitude errors, as measured by lnMAG, are sensitive to anatomical errors only. Consequently, magnitudes from a template model or simplified models cannot be trusted. Based on the results presented here, we suggest to use a forward model as realistic as possible. The availability of pipelines able to generate realistic geometrical descriptions of the human head have so far been limited. SimNIBS makes it easy to automatically generate models and the computational cost of doing so is not greater than running a FreeSurfer pipeline (as required when using MNE-Python) or the DUNEuro solver (used in the Fieldtrip pipeline). See Appendix A for an example.

If digitized electrode positions are unavailable, we suggest to use a template created by measuring the positions on a realistically shaped head template. Alternatively, average positions from a previous study using the same cap could be used. The template described here will be included in SimNIBS and we plan to add other caps in the future. Perhaps somewhat surprisingly, using a geometrical model based on template anatomy did not result in larger errors than coarse anatomical modeling of the individual anatomy. This suggests that if individual anatomical information is lacking then it might still be possible to generate a usable forward model based on standard anatomy. Even though we did not evaluate the impact of the observed effects on source localization accuracy, previous studies suggest that our main findings will translate to source localization although this association will likely be modulated by the noise level in the data and the particular choice of inverse solver.

## Acknowledgements

JDN is supported by a PhD stipend from Sino-Danish Center for Education and Research. OP was supported by the Lundbeck foundation (grant R360-2021-395). AT was supported by the Lundbeck foundation (grants R244-2017-196 and R313-2019-622) and the NIH (grant 1RF1MH117428-01A1). The authors would like to thank Fan Wang (Chinese Academy of Sciences) for helpful discussions regarding electrophysiological data acquisition and modeling.

## Data Availability

The data used in this study has been presented previously and we refer to the original articles for statements about data availability. The data used to compare the different forward modeling pipelines was presented by Farcito et al. (2019). The data used to compare electrode specification strategies was presented by Madsen et al. (2019) and Karabanov et al. (2021).

**Listing 1.**
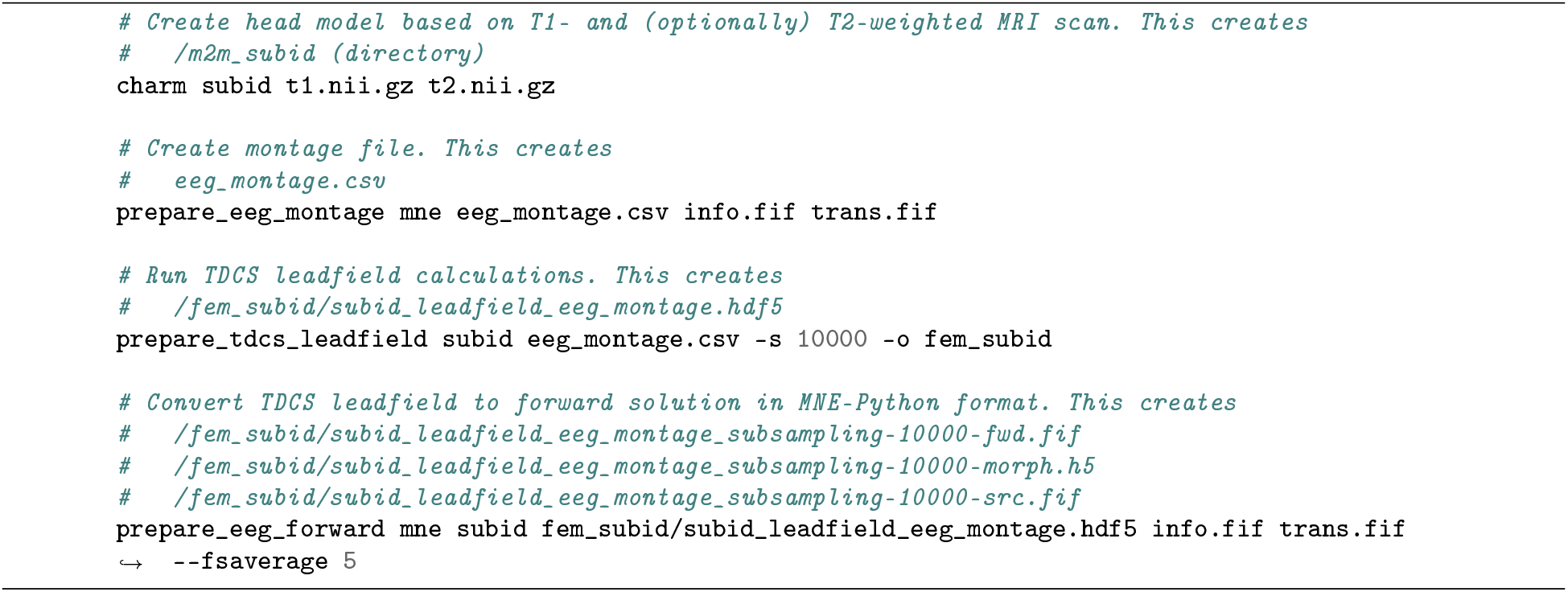
Generating an EEG forward solution for MNE-Python with SimNIBS. `info.fif` can be a Raw, Epochs, Evoked or Info object. `trans.fif` is a Trans object containing an affine mapping between head coordinates and subject MRI coordinates. `-fwd.fif, -morph.fif`, and `-src.fif` are standard MNE-Python objects containing the forward solution, the sparse matrix to morph to *fsaverage* space, and the source space definition, respectively.

## Appendix A. Generating EEG Forward Solutions With SimNIBS

Here we show how to generate EEG forward solutions with SimNIBS which can be used with MNE-Python (listing 1) or FieldTrip (listing 2). In both examples, the central gray matter surface of each hemisphere is subsampled to 10,000 nodes per hemisphere and a mapping to *fsaverage5* (which contains 10,242 nodes per hemisphere) is constructed. In both cases, the procedure is the same but the output format is different: when exporting to MNE-Python format (listing 1), the forward solution is in the head coordinate system used by MNE-Python whereas when exporting to FieldTrip (listing 2), the forward solution is in subject MRI coordinate system.

---

**Listing 2.**
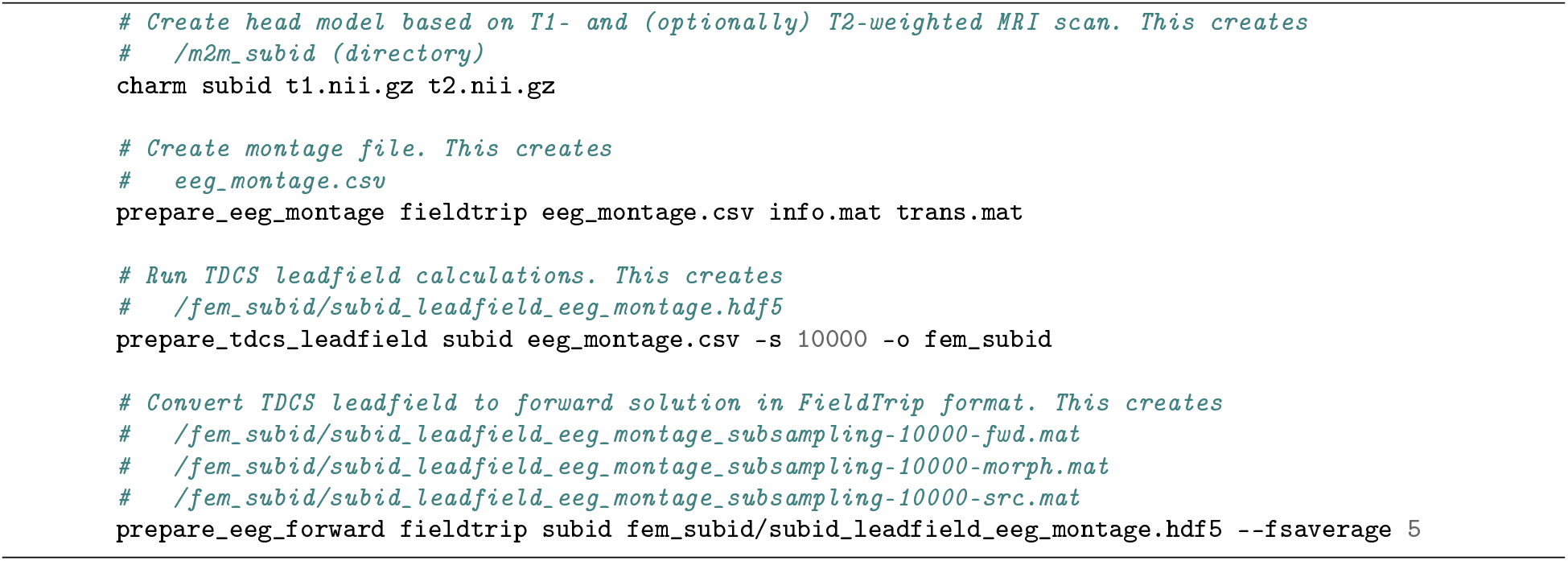
Generating an EEG forward solution for FieldTrip with SimNIBS. `info.mat` contains information about the electrodes and `trans.mat` contains an affine mapping between the coordinate system of the electrodes and MRI space. `-fwd.mat` contains the forward solution, `-morph.mat` is a sparse matrix which can be used to “morph” from subject space to *fsaverage* space, and `-src.mat` contains the source space definition.

## Footnotes

1 Efficient ways of solving the system, e.g., the fast multipole method, exist allowing for high resolution BEM models as well.

2 Instead of a current density, we simply consider a point dipole with unit Am and no associated (or infinitesimally small) volume and thus we have dropped the *δV* term.

3 Using point electrodes (as is common in EEG forward calculations) did not have a strong influence and so here we present the results for meshed electrodes which is the default approach in SimNIBS.

4 White matter, gray matter, CSF, spongy bone, compact bone, skin, eyes, veins, muscle.

5 Upon visual inspection, many sulci were closed in the manual segmentations, hence these were “corrected” using FreeSurfer by (1) enforcing gray matter within the cortical gray matter mask from FreeSurfer, (2) relabeling any cortical gray matter voxels outside of this mask which overlapped with the white matter mask from FreeSurfer to white matter, and (3) relabeling any unlabeled voxels to CSF. This seemed to work well for opening sulci, however, at the price of slightly reduced accuracy around cerebellum.

6 MNE-Python also provides a tool to create surfaces from a T1-weighted image and a sequence of fast low angle shot (FLASH) images, however, since FLASH was not available for this dataset we decided to only use the T1-weighted image (we tried using the T2-weighted image instead but with disappointing results).

7 We used the master branch from the GitHub repository (https://github.com/fieldtrip) at commit 4b50f71105f6bb250018b815bb628e6f3e9de2b6 (from 22.11.2021).

8 For FieldTrip-SPM, we specified conductivities as defined in the tutorial https://www.fieldtriptoolbox.org/tutorial/headmodeleegfem.

9 When sources were excluded, smoothing was employed to ensure that all nodes in *fsaverage* space were covered. To avoid unnecessary smoothing, this was only done for the nodes where it was required.

10 The cap used here has an equidistant M10 layout and includes two additional channels in the temporal area. Here we use the manufacturer layout for this cap shipped with MNE-Python augmented by these two channels. To adhere to the equidistant properties of the layout, we defined the angles (theta, phi) of these electrodes as (−138, −22) and (138, 22).

11 The conductivity of compact and spongy bone were both set to that of bone in the FieldTrip-SPM model and thus a minor geometrical simplification of the reference was introduced.

## Notes

### Competing Interest Statement

The authors have declared no competing interest.

