## Supplementary material for "Evaluating the Influence of Anatomical Accuracy and Electrode Positions on EEG Forward Solutions"

### S1 SimNIBS Validation

Figure S1.1 shows that the numerical accuracy of finite element method (FEM) generally deteriorates as sources get closer to the sensors in a realistically shaped three-compartment model. The boundary element method (BEM) solution from MNE-Python was used as reference. We also compare with a BEM solution from FieldTrip to show how this is more numerically accurate for high-eccentricity sources (at these resolutions). Figures S1.2 and S1.3 show error as a function of eccentricity in a realistically shaped five-compartment model, however, in this case the resulting impact of eccentricity is less clear. A high resolution FEM model from SimNIBS was used as reference.

### S2 Additional Illustrations of Volume Conductor Geometry

To support the claims in the main text about the general differences between the volume conductor models from the different pipelines in the anatomy study, we show a few more examples in figures S2.4 to S2.7.

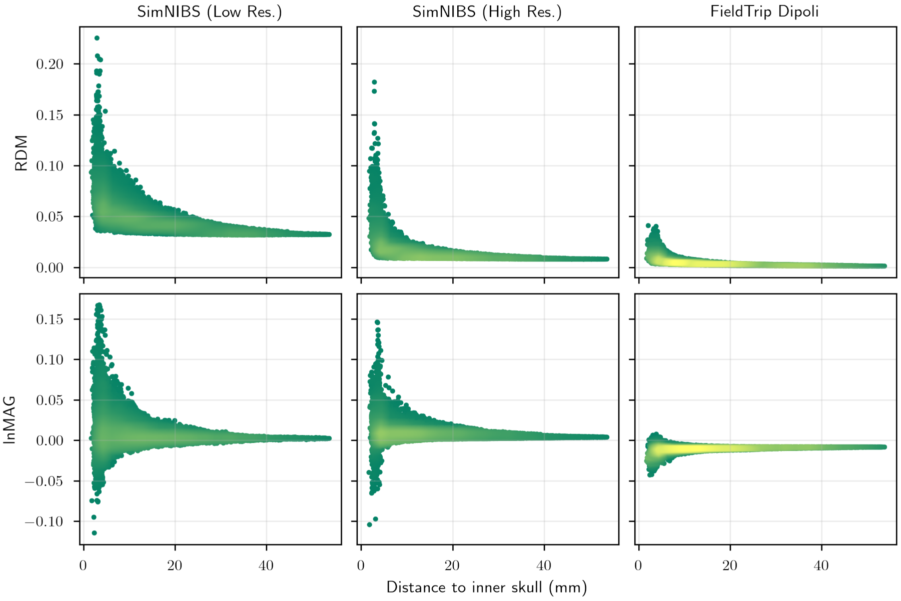


**Figure S1.1.** Average error (relative difference measure (RDM) or logarithm of the magnitude error (lnMAG)) per source location as a function of eccentricity for a high and low resolution FEM model and a BEM model in a realistically shaped three-compartment model. The BEM solution from MNE-Python was used as a reference.

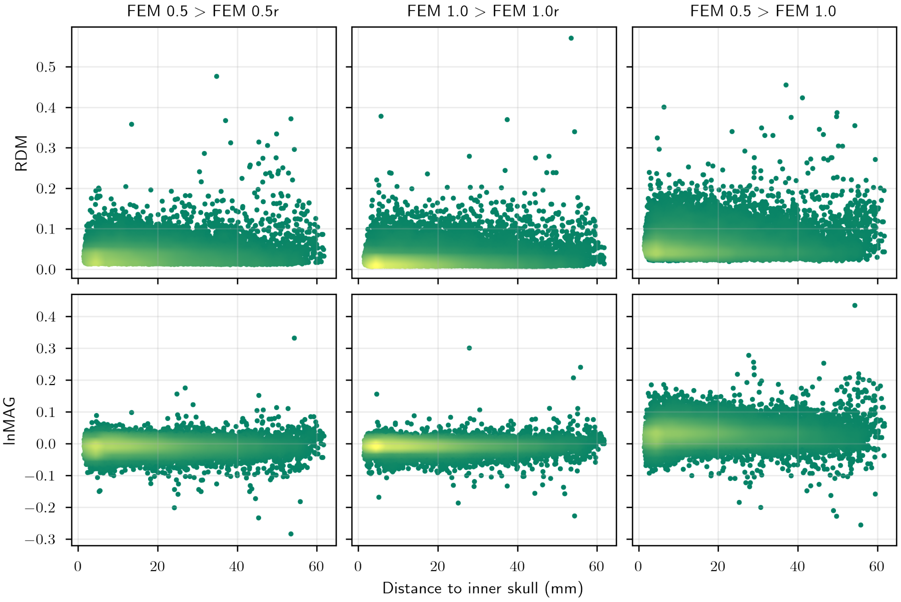


**Figure S1.2.** Average error (RDM or lnMAG) per source location as a function of eccentricity for the FEM models.

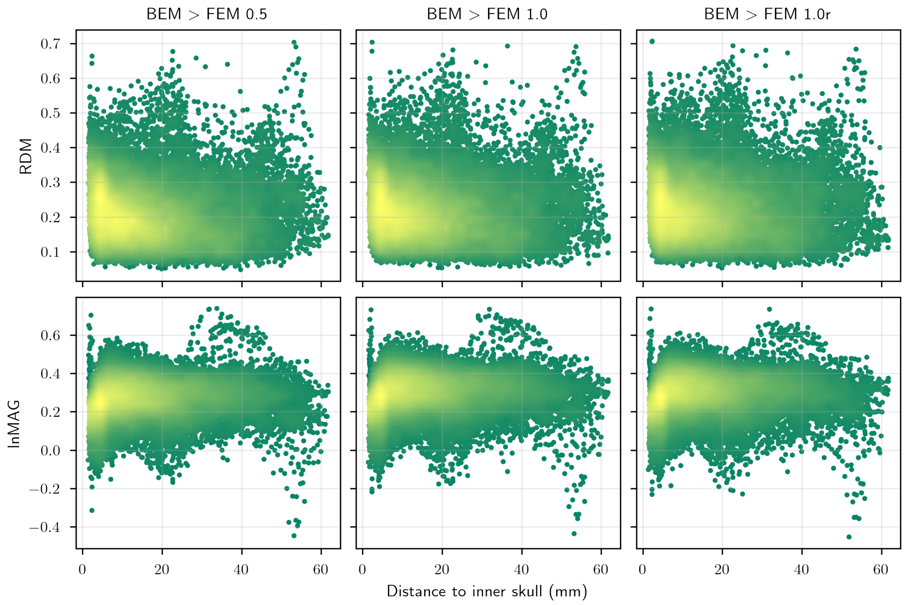


**Figure S1.3.** Average error (RDM or lnMAG) per source location as a function of eccentricity for the BEM model.

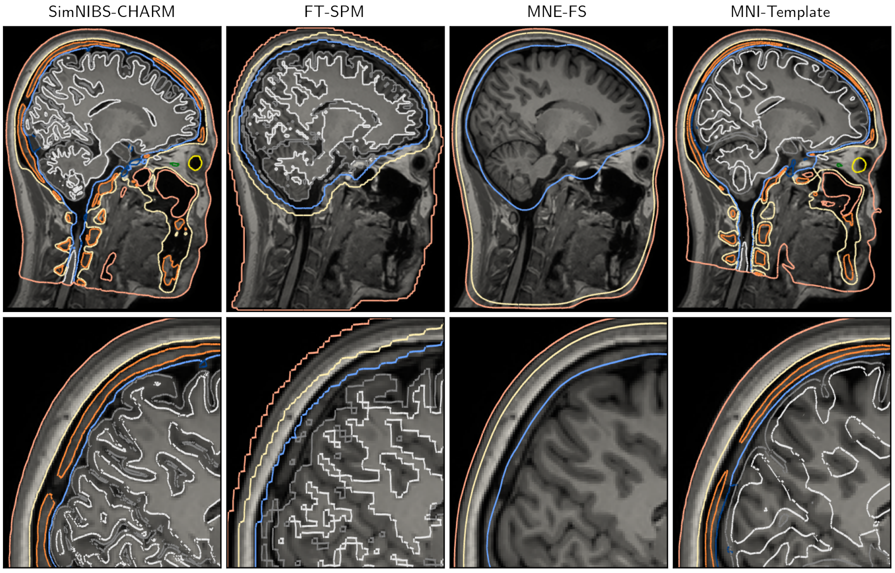


**Figure S2.4.** Tissue compartments as estimated by each of the pipelines in the anatomy study for subject 3.

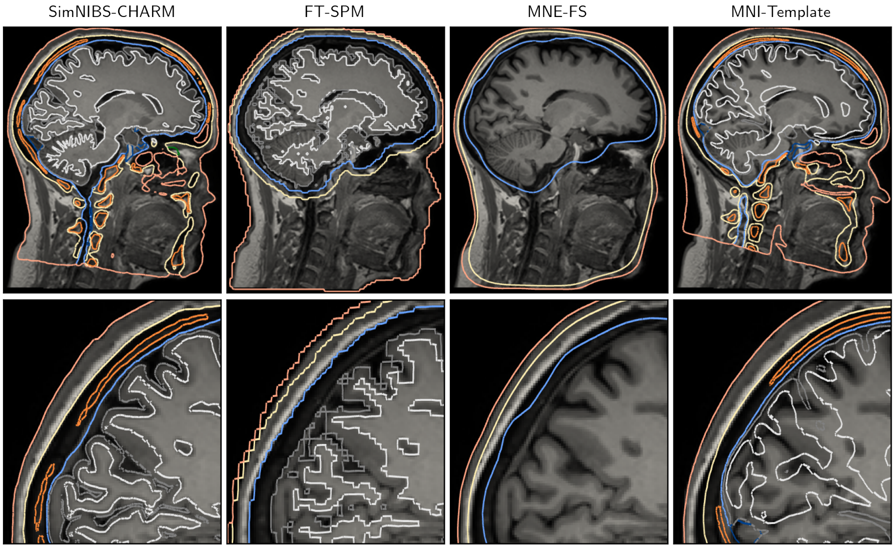


**Figure S2.5.** Tissue compartments as estimated by each of the pipelines in the anatomy study for subject 4.

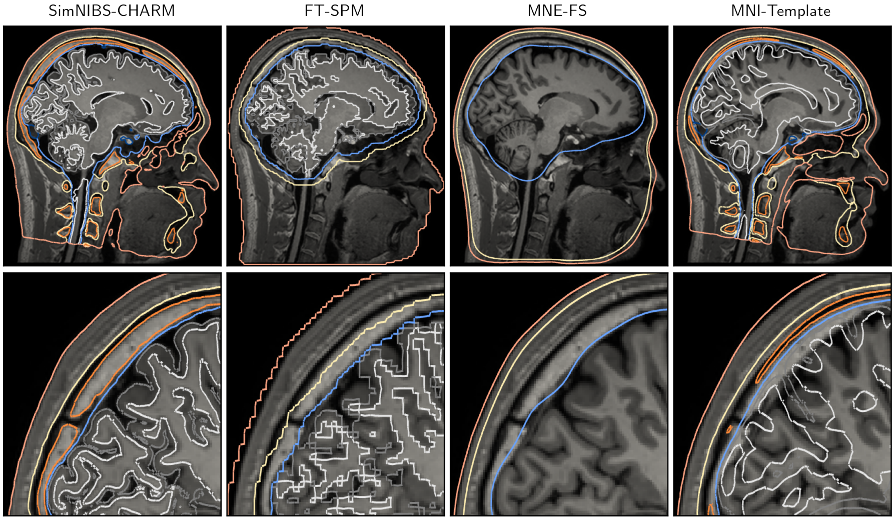


**Figure S2.6.** Tissue compartments as estimated by each of the pipelines in the anatomy study for subject 5.

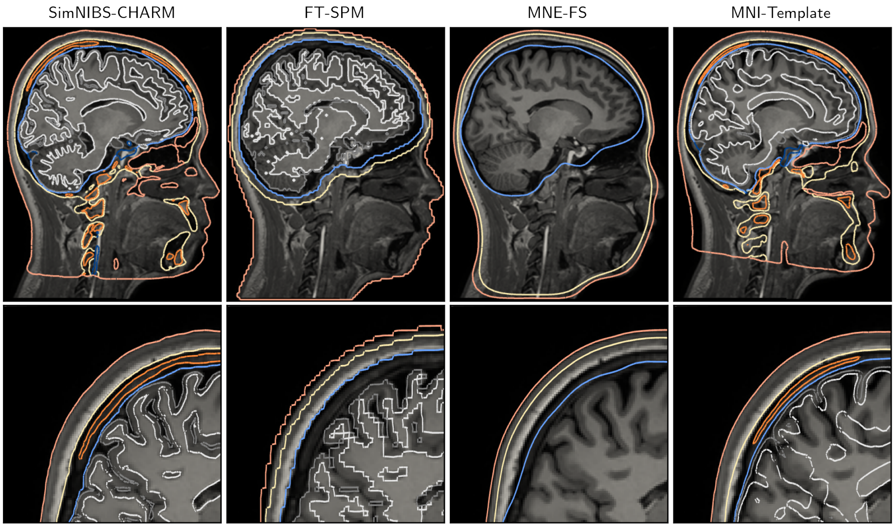


**Figure S2.7.** Tissue compartments as estimated by each of the pipelines in the anatomy study for subject 6.
